# Mucoidy, a general mechanism for maintaining lytic phage in populations of bacteria

**DOI:** 10.1101/775056

**Authors:** Waqas Chaudhry, Esther Lee, Andrew Worthy, Zoe Weiss, Marcin Grabowicz, Nicole Vega, Bruce Levin

## Abstract

We present evidence that phage resistance resulting from overproduction of exopolysaccharides, mucoidy, provides a general answer to the longstanding question of how lytic viruses are maintained in populations dominated by bacteria upon which they cannot replicate. In serial transfer culture, populations of mucoid *E. coli* MG1655 that are resistant to lytic phages with different receptors, and thereby requiring independent mutations for surface resistance, are capable of maintaining these phages with little effect on their total density. Based on the results of our analysis of a mathematical model, we postulate that the maintenance of phage in populations dominated by mucoid cells can be attributed primarily to high rates of transition from the resistant mucoid states to susceptible non-mucoid states. Our tests with both population dynamic and single cell experiments as well as DNA sequence analysis are consistent with this hypothesis. We discuss reasons for the generalized resistance of these mucoid *E. coli*, and the genetic and molecular mechanisms responsible for the high rate of transition from mucoid to sensitive states responsible for the maintenance of lytic phage in mucoid populations of *E. coli.*

## Introduction

Bacteriophages are natural predators (parasitoids) of bacteria and are present in natural microbial communities across environments, from marine and aquatic systems (Chibani-Chennoufi et al. 2004) to the intestine of the mammalian host (Abeles and Pride 2014). By mutation, bacteria can become resistant (Campbell 1961) or, in the case of CRISPR-Cas, immune to these viruses (Barrangou et al. 2007). Most commonly these resistant or immune cells ascend to dominate the bacterial population, but the phage continue to be maintained (Horne 1970; Chao, Levin, and Stewart 1977; Wei, Ocampo, and Levin 2010; Gomez and Buckling 2011; Chaudhry et al. 2018). How virulent phage are maintained in populations of bacteria dominated by cells upon which they cannot replicate is a longstanding question that has been addressed theoretically as well as experimentally (Campbell 1961; Levin, Stewart, and Chao 1977; Jover, Cortez, and Weitz 2013; Jover, Romberg, and Weitz 2016; Weissman et al. 2017). Thanks to concerns about the factors responsible for maintaining and stabilizing the structure of bacterial communities, microbiomes, if you prefer (Reyes et al. 2013; Manrique et al. 2016) and the resurrection of interest in phage therapy (Bull, Levin, and Molineux 2019; Schmidt 2019; Cafora et al. 2019; Forti et al. 2018), this question - and more broadly, studies of the population and evolutionary dynamics of the interactions between bacteria and phage - have become an increasingly important and relevant avenue of research.

Those who work with *E. coli* and other Gram negative bacteria occasionally see glossy, raised, opaque “mucoid” colonies on agar that can be attributed to contamination with bacteriophage. By providing a physical barrier between the cell surface and the would-be infecting phage the overproduction of the exopolysaccharide responsible for the mucoid phenotype confers resistance to the phage (Scholl, Adhya, and Merril 2005; Bernheimer and Tiraby 1976; van der Ley, de Graaff, and Tommassen 1986; Labrie, Samson, and Moineau 2010; Wilkinson and Holmes 1979). While this resistance may not be as complete as that generated by the modification of the phage receptors (Harrison et al. 2015) and may engender a substantial fitness cost (Wielgoss et al. 2015), it is sufficient to select for mucoidy in populations of bacteria exposed to lytic phage (Mizoguchi et al. 2003; Scanlan and Buckling 2012; Wielgoss et al. 2015). Moreover, and most importantly from an ecological perspective, there is evidence that the resistance provided by mucoidy is general rather than specific, like the resistance generated by modification of the phage receptors; mucoid bacteria are resistant to multiple lytic phages (Wielgoss et al. 2015).

These observations raise several questions about the population dynamics and ecology of mucoid-based resistance. Under what conditions will mucoidy be selected for over surface (envelope) or other resistance or immune mechanisms (Labrie, Samson, and Moineau 2010)? How might the ecology of phage-bacterial systems differ when resistance is conferred by mucoidy rather than envelope resistance? Why and how, in the absence of phage, do mucoid-colony producing bacteria revert to states that produce non-mucoid colonies? Most importantly, what is the contribution of mucoidy to the coexistence of bacteria and lytic phage?

To address and answer these questions, we studied the population dynamics of mucoid colony producing *E. coli* K12 strain MG1655 following exposure to different virulent (lytic) bacteriophage, T3, T4, T5, T7 and □^VIR^. We present evidence that mucoidy provides a sufficient answer to the question of how lytic phage are maintained following the evolution of resistance and a general mechanism for the stable co-existence of bacteria with lytic phage. Based on the results of our analysis of mathematical/computer simulation models, we postulate that lytic phages can stably persist on populations of mucoid *E. coli* by replicating on minority populations of sensitive bacteria that are maintained primarily if not exclusively by a high rate of transition from “resistant” mucoid to sensitive non-mucoid colony types. Using a combination of computer simulations, batch culture experiments, single-cell microscopy, and DNA sequence analysis we present evidence in support of this “leaky resistance” (Chaudhry et al. 2018; Weissman et al. 2017; Delbruck 1946) hypothesis for the maintenance of lytic phage in populations of mucoid bacteria.

## Materials and Methods

### Bacteria, phage and growth media

Strains used in these experiments are shown in Table 1. Bacterial cultures were grown at 37°C either in M9 media [M9 salts (248510, Difco)] supplemented with 0.4% glucose (6363-53-7, Fisher Scientific), 1 mM MgSO_4_ (Sigma Aldrich), 0.1 mM CaCl_2_ (JT Baker), and 0.2% Thiamine B1 (Sigma Aldrich)] or LB broth [MgSO_4_ 2.5g/L, tryptone (Fisher Bioreagent 10g/L, yeast extract (Bacto) 5g/L, sodium chloride (Fisher Chemical) 10g/L]. All *E. coli* strains used in our experiments were derivatives of the parent strain K12 MG1655. Rcs pathway knockout strains were constructed by the method described in (Baba et al. 2006). Gain-of-function point mutant for mucoid phenotype (*rcsC137*) (Gottesman, Trisler, and Torres-Cabassa 1985) was introduced via co-transduction with *ompC*::Tn*5* (this *ompC* allele does not cause phage resistance and is a neutral marker in our experiments).

**Table 1.**
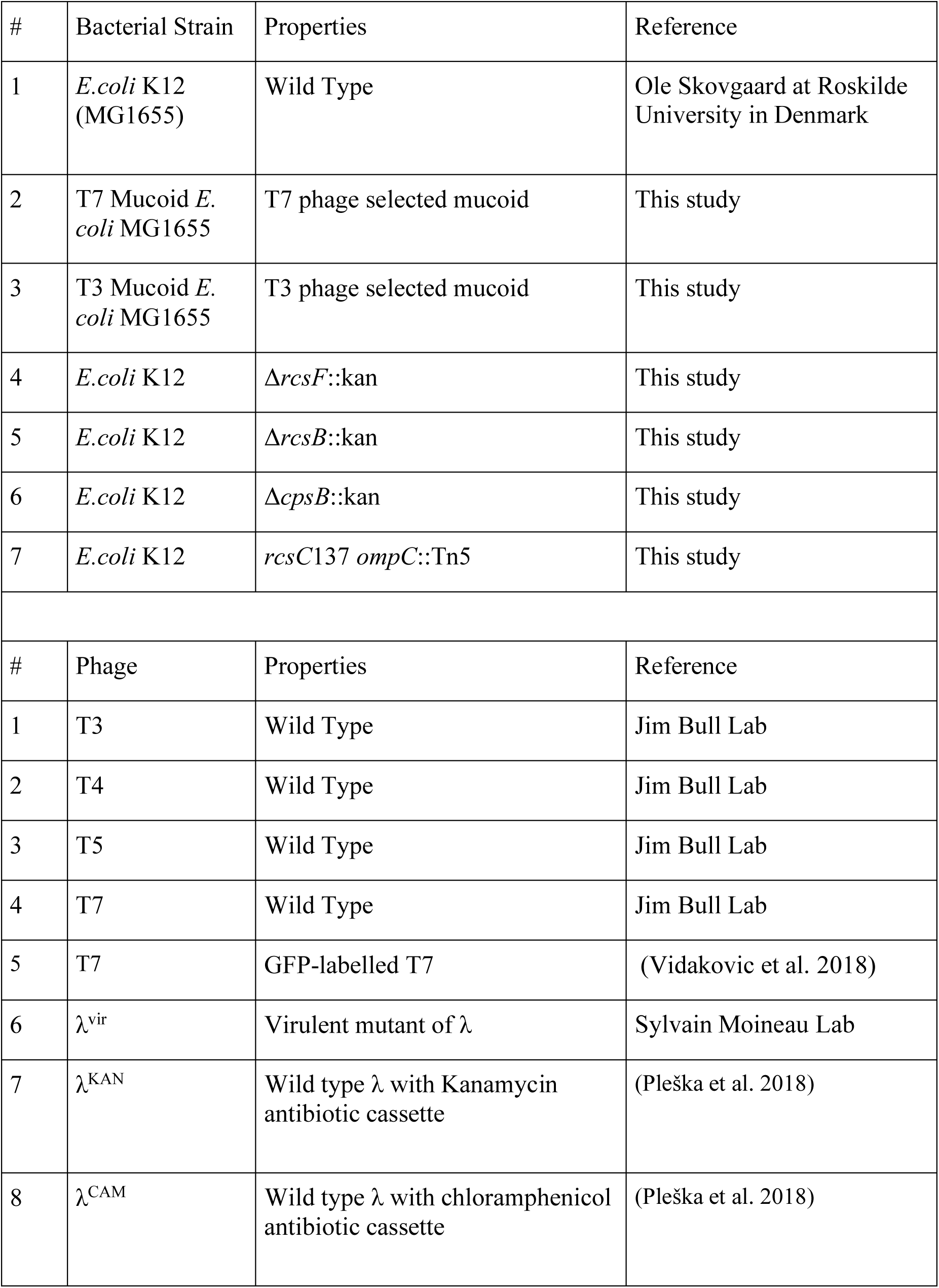
Bacterial strains and phage used in this study

Phage lysates were prepared from single plaques at 37°C in LB medium alongside wild-type *E. coli* MG1655. Chloroform was added to the lysates and the lysates were centrifuged to remove any remaining bacterial cells.

The λ^VIR^ strain used in these experiments was obtained from Sylvain Moineau (Laval University, Quebec City, Canada) The construction of temperate phage λ^KAN^ and λ^CAM^ is described in (Pleška et al. 2018).

### Sampling bacterial and phage densities

Bacteria and phage densities were estimated by serial dilutions in 0.85% NaCl solution followed by plating. The total density of bacteria was estimated on LB (1.6%) agar plates. To estimate the densities of λ^KAN^ or λ^CAM^ lysogens, cultures were plated on LB (1.6%) agar with 25 μg/mL kanamycin (AppliChem Lot# 1P0000874) or chloramphenicol (Merck Lot# 2430B61), respectively.

To estimate the densities of free phage, chloroform was added to suspensions before serial dilutions. These suspensions were mixed with 0.1 mL of overnight LB grown cultures of wild-type MG1655 (about 5×10^8^ cells per mL) in 3 mL of LB soft (0.65%) agar and poured onto semi-hard (1%) LB agar plates.

### Serial Transfer

All serial transfer experiments were carried out in 10-mL cultures in 50 ml Erlenmeyer flasks grown at 37°C with vigorous shaking. The cultures were initiated by 1:100 dilution from 10-mL overnight cultures grown from single colonies. Phage were added to these cultures to reach the initial density of approximately 10^6^ PFU/mL. At the end of each transfer, 100 μL of each culture was transferred into flasks with fresh medium (1:100 dilution). Simultaneously, 100 μL samples were taken for estimating the densities of colony forming units (CFU) and plaque forming units (PFU), by serial dilution and plating on solid agar, with selection as needed as described above.

### Bacteriophage and bacteria parameter determination

The parameters critical for the interaction of λ phages and *E. coli* MG1655 used in this study were estimated in independent experiments in LB medium. The maximum growth rate of *E. coli* MG1655 was measured by Bioscreen as described in (Concepción-Acevedo et al. 2015). Phage burst sizes (β) were estimated with one-step growth experiments (Ellis and Delbrück 1939) in a manner similar to (Abedon, Herschler, and Stopar 2001). Adsorption of λ to *E. coli* was estimated as described in (Ellis and Delbrück 1939). The procedure for estimating the probability of lysogeny and the rate of spontaneous lysogen induction are presented in (Pleška et al. 2018).

### Short term dynamics experiments

Overnight cultures of *E. coli* MG1655 were grown in LB media. The cultures were diluted in a 1:1000 ratio on 20 mL of medium in 100 mL flasks to which λ_KAN_ or λ_CAM_ phage was added to an initial density of 1×10^7^ PFU/mL. Short term dynamics were conducted culturing the mixture at 37°C in shaking conditions. Total bacteria count was conducted by sampling 100 μL at regular intervals and plating dilutions on LB agar plates. Lysogen counts were obtained by sampling 100 μL and plating dilutions on LB agar with 25 μg/mL kanamycin or chloramphenicol. Phage counts were obtained by sampling 1 mL at regular intervals, adding chloroform to kill bacteria, and plating for PFU quantification as previously described.

### Resistance Testing

We probed for resistance using two different assays: one in liquid and another using the double layer soft agar technique.

Liquid culture: Colonies from the bacteria plates were streaked thrice to ensure removal of phage. Colonies from the streaked plates were used to establish overnight cultures in 2 mL LB broth incubated at 37°C in a shaker. The overnight cultures were diluted 1/100 and ∼10^5^ PFU/ml of phage were added to the cultures. After 24 hours of incubation, free phage in these cultures were measured. Those unable to support the phage replication were considered as resistant (Wei, Ocampo, and Levin 2010).

Phage agar: Phage plates were made as previously described, using 0.1 mL of overnight LB grown cultures of the isolates being tested. Dense phage stocks (10^8^-10^9^ PFU/mL) were spotted onto the agar surface; and, susceptibility was scored according to the presence of visible plaques.

### Single Cell Microscopy

Overnight cultures of WT and mucoid *E.coli* MG1655 carrying plasmid pBAD18Cm::P_rprA_-mCherry (reporter for induction of the Rcs pathway (Konovalova et al. 2018) were grown in LB media. These cultures were diluted 1/1000 in fresh LB and incubated for 30 minutes with vigorous shaking (300 RPM) at 37°C. We then added GFP-labelled T7 phage (Vidakovic et al. 2018) at 0.1 MOI and returned to incubation in a stationary incubator at 37°C. The phage free control and phage infected culture slides were sampled at 15-30 minute intervals and observed on an inverted fluorescent microscope (Leica DMi8 with LasX Premium software). Images are an overlay of DIC, TRITC, and GFP channels, acquired using a 40X objective (total magnification 400X). Quantification was performed using Image J. Object thresholds were established using DIC images, and particle analysis was performed on all particles exceeding 0.1 µM^2^ area. Resulting plots show mean pixel intensity in each fluorescent channel for all identified objects.

### DNA sequencing

Genomic DNA was isolated using Promega Wizard genome DNA purification kit (Catalog Number PAA1120) according to manufacturer instructions. Illumina pair-ended libraries were prepared from genomic DNA and sequenced by Microbial Genome Sequencing Center Pittsburg (MiGS). Sequencing reads were assembled into contigs using SPAdes Assembler (version 3.14) (Nurk et al. 2013). Contigs from parental strains, mucoid mutants, and non-mucoid revertants were aligned with the MG1655 reference sequence (GenBank ID U00096.2) and analyzed for mutations using progressiveMauve (Darling, Mau, and Perna 2010).

## Results

### I Selection for and the population dynamics of mucoid *E. coli* in phage-infected cultures

First, we sought to determine whether lytic phages varied in their ability to select for mucoidy in *E. coli* MG1655, and whether these phage-selected mucoids differed from one another in their interactions with phage. For this, wild type cultures of *E. coli* MG1655 were exposed to T3, T4, T5, or T7 phage and serially transferred over ten passages (Figure 1). For cultures initiated with T3 or T4, the phage were maintained over all ten transfers and the dominant population of bacteria isolated at the end of the final transfer was mucoid (Figure 1A and 1B). All replicates of the experiments using T5 resulted in the extinction of the phage following the emergence of envelope resistance (Figure 1C). For all our T5-resistant isolates, we confirmed that envelope resistance mapped to the *fhuA* gene which encodes the T5 receptor. We expect that all these isolates carry *fhuA* null mutations since such mutations are well known to confer T5 resistance (Dykhuizen and Dean 2004). Indeed, whole-genome sequencing of one isolate confirmed an IS*1* insertion in *fhuA* that disrupts the gene and prevents receptor production. For T7, the phage were maintained in one of the three populations, where mucoid cells were the dominant population of bacteria. In the remaining two T7 cultures, neither mucoid nor resistant mutants were detected and bacteria and phage were rapidly driven to extinction (Figure 1D).

**Figure 1:**
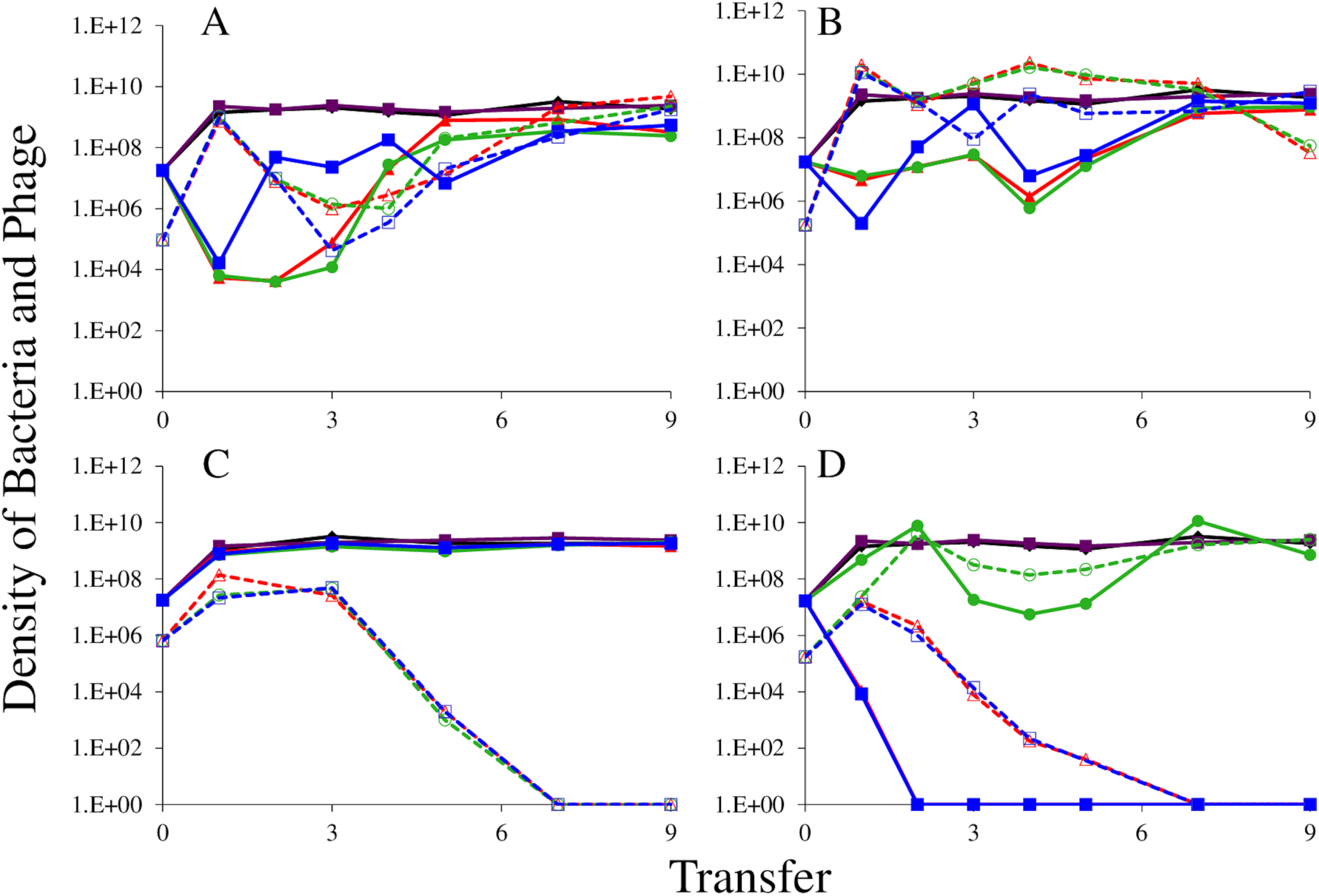
The population dynamics of T3 (A), T4 (B), T5 (C) and T7 (D) phage with wild-type *E. coli* MG1655. Cultures were diluted 1:100 into fresh LB medium every 24 hours; three replicates were performed per condition. Solid lines show bacterial densities in each replicate (red/green/blue); dotted lines show the phage densities in the corresponding cultures. Phage-free cultures (black and purple lines) are shown as controls, to indicate resource-limited population size for the bacteria.

Second, we explored the capacity of these phages to be maintained in populations of mucoid bacteria. For the first of these serial transfer experiments, we challenged mucoids generated by selection with the phage T7 with each of T3, T4, T5, or T7 phage. As seen in Figure 2, all four phage were maintained on T7-derived mucoids, and the densities of viable bacteria were high, similar to that of the phage-free controls. Stated another way, resources rather than phage limited the densities of these bacterial populations. Moreover, as can be seen in Figure S1, this result obtains in minimal medium as well as the LB used for the Figure 1 experiment. In the minimal medium experiments, however, mutants with envelope resistance arose in mucoid cultures treated with phage T4 and T5.

**Figure 2:**
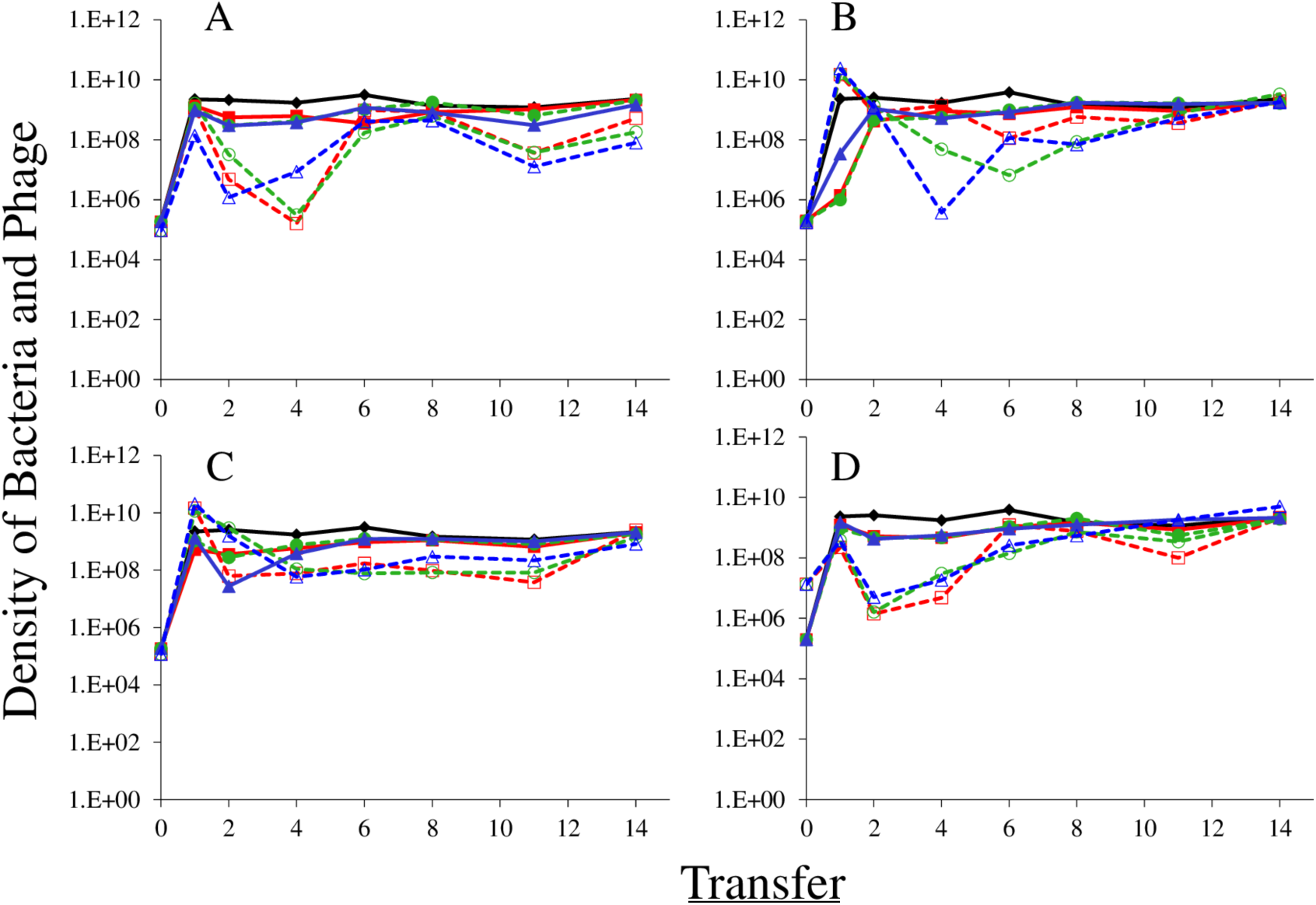
The population dynamics of phage T3 (A), T4 (B), T5(C) and T7 (D) phage with T7-selected mucoid *E. coli* MG1655 as the host strain in LB. Three independent cultures were diluted 1:100 into fresh LB media every 24 hours, and CFU/PFU estimates of the densities of bacteria and phage were made by serial dilution plating. Solid lines show bacterial densities in each replicate (red/green/blue); dotted lines show the phage densities in the corresponding cultures. Phage-free cultures (black and purple lines) are shown as controls, to indicate resource-limited population density for the bacteria. Phage-free controls lost the mucoid phenotype entirely by the end of the 4^th^ serial transfer. At the end of the 15^th^ serial transfer, three colonies from each phage-infected culture were streaked at least three times to get rid of phage, yielding non-mucoid revertants that were again sensitive to the relevant phage.

These results support the initial assumption that phage-selected mucoidy in *E. coli* provides broad resistance to virulent phage and indicate that co-existence of large populations of virulent phage and mucoid *E. coli* is a general phenomenon that does not depend on the phage initially used for selection. These results indicate that the capacity to select for mucoidy and/or the rate of mutation to mucoidy varied among the phages. These results also suggest that mucoidy can prevail over envelope resistance, but may not do so in all cases, depending on the environment and on the lytic phage used.

### II Mechanism of phage-selected mucoidy

Next, we investigated the ability of defined mutations in the *E. coli* colanic acid synthesis pathway to recapitulate the observed ecological interactions between bacteria and phage, specifically the high-level maintenance of phage on apparently resistant hosts. The “regulator of capsule synthesis” (Rcs) phosphorelay system regulates colanic acid capsule production in *E. coli* (Clarke 2010). The Rcs phosphorelay is complex, but at its core: RcsF is a sensory lipoprotein that causes activation of the RcsC sensor kinase in an environment-dependent manner; RcsC causes phosphorylation of the RcsB transcription factor; phosphor-RcsB is a positive regulator of the colanic acid biosynthetic operon at the *cpsB* promoter. Overproduction of colanic acid via this pathway results in the distinctive mucoid colony phenotype, and mucoid strains featuring mutations in this pathway are selected for by exposure to virulent phage in *E. coli* MG1655 (Wielgoss et al. 2015).

To understand the dynamics of mucoid selection and the role of mucoidy in phage maintenance, we sought to determine whether EPS-impaired strains, which do not produce mucoid colonies, would be able to maintain phage populations during serial passage. To this end, we tested two gene knockout strains in the Rcs signaling pathway (Δ*rcsB* and Δ*rcsF*) as well as one gene knockout strain in the EPS operon (Δ*cpsB*), all of which are impaired in capsule synthesis. We also tested a strain (*rcsC137 ompC::Tn5*) that has an activating A904V substitution in RcsC and is therefore constitutively mucoid (Stout 1996), to determine if the simple induction of mucoidy via a known mechanism was sufficient to allow maintenance of phage. We found that the *rcsC*137 constitutive-mucoid mutant was able to maintain large populations of phage and bacteria for all phage tested, while phage exposure in loss-of-function mutants impaired in mucoidy (Δ*rcsB*, Δ*rcsF*, Δ*cpsB*) resulted in extinction of both phage and bacterial populations when T3 phage were used (Fig. 3). Exposure to T4 or T7 phage resulted in bacterial population collapse in some conditions but not others, largely dependent on whether evolution of envelope resistance occurred; in the Δ*rcsF* background, mucoid colonies were observed in the last 5^th^ serial transfer cultures of T4, presumably downstream of a mutation compensating for the mucoidy defect in the knockout. For reasons that are not yet clear, T5 phage were maintained on all host backgrounds regardless of mucoidy, despite the fact that evolution of envelope resistance generally leads to extinction of this phage (Fig. 1).

**Figure 3.**
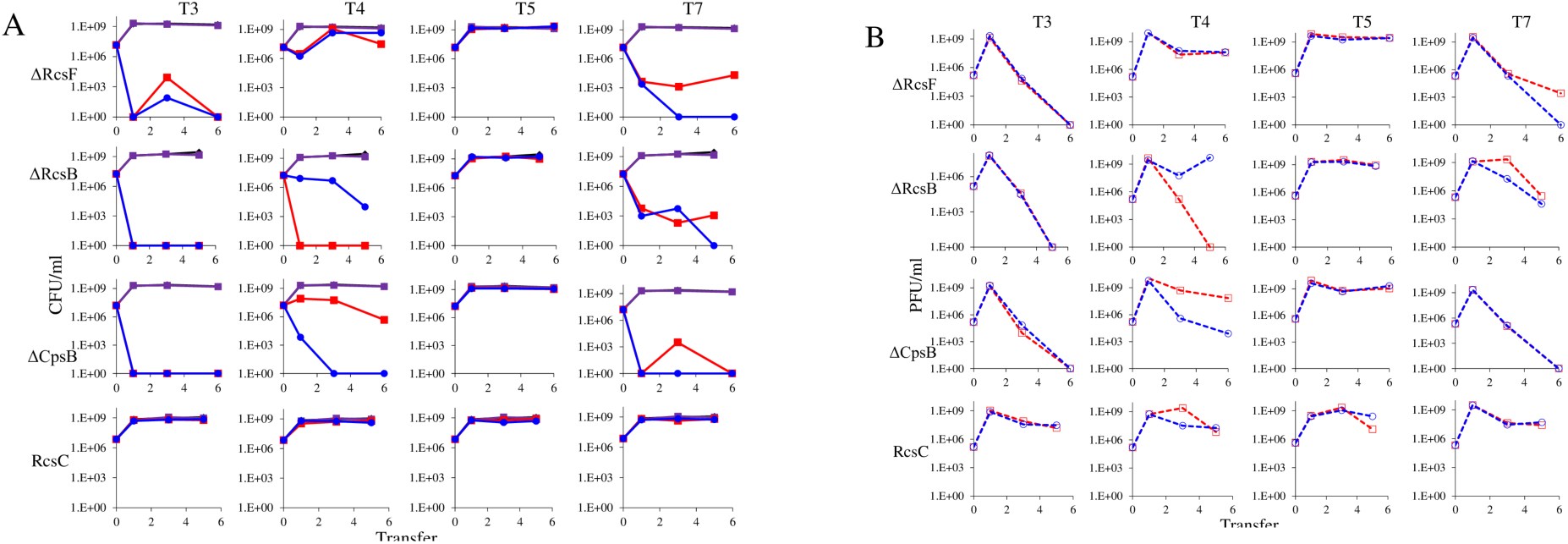
Constitutive presence or absence of mucoidy affects the dynamics of *E. coli*-phage interactions. T3, T4, T5, or T7 phages were co-cultured with genetically modified strains of *E. coli* MG1655 with known gain or loss of function in the Rcs pathway. Strains used were: Δ*rcsF*::kan (pathway loss-of-function knockout of the outer membrane Rcs input), Δ*rcsB*::kan (pathway loss-of-function knockout of the RcsB transcription factor), Δ*cpsB*::kan (pathway loss-of-function knockout of biosynthesis of exopolysaccharide colonic acid), and *rcsC*137 *ompC*::Tn5 (gain-of-function point mutant that is a defined activating mutation of mucoidy). Cultures were diluted 1:100 into fresh LB medium every 24 hours; two replicates were performed per condition. Phage-free cultures are shown as controls (black and purple lines), to indicate resource-limited population size for the bacteria. Solid red and blue lines show bacterial densities (A), while dotted red and blue (B) lines show the phage densities in the corresponding cultures.

### III A hypothesis for the maintenance of phage in mucoid populations of bacteria

We now consider the population dynamics of lytic phage and mucoid hosts to address two questions. How are the phage maintained in mucoid populations of bacteria? Under what conditions will envelope resistant mutant bacteria emerge and replace the mucoid population?

We postulate that the maintenance of high densities of phage in populations of mucoid bacteria can be attributed to a substantial density of sensitive cells generated by a high rate of transition from the mucoid to non-mucoid states, a mechanism that has been called “leaky” resistance (Chaudhry et al. 2018). To illustrate how this mechanism can account for the observed maintenance of high densities of phage with little effect on the density of bacteria, and for the common failure to see the ascent of mutants with surface resistance, we use a simple mass-action model of the population dynamics of bacteria and phage (Fig.4, Equations 1-5). In this model, there are three populations of bacteria – sensitive (S), mucoid (M), and resistant (N_R_), and a single population of lytic phage (P), where S, M, N_R_, and P are the densities, cells or particles per ml, as well as the designations of these populations.

**Figure 4:**
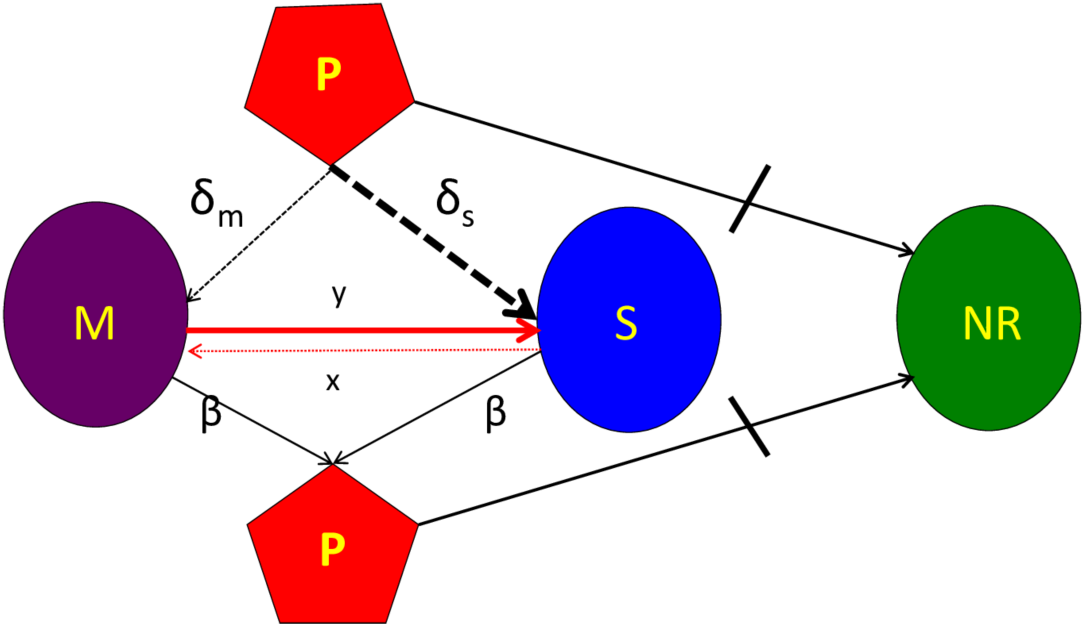
Diagram of the mass-action model, showing transitions between states. Bacteria may be mucoid (M), sensitive (S) or envelope-resistant (NR). Mucoid and sensitive cells adsorb phage (P), but resistant cells do not (as shown by the stopped lane). See the text and methods for full definitions of the parameters, variables, and assumptions behind the construction of this model. The resource, r, is not considered in this diagram, but is included in the numerical solutions to these equations, which simulates a serial transfer mode of population maintenance.

The bacteria grow at maximum rates *v*_*S*_, *v*_*M*_ and *v*_*NR*_ (cell^-1^ hr^-1^) respectively (S2 Fig), with the realized growth rates dependent on the concentration of the shared limiting resource (r, µg/mL) via a hyperbolic function ψ(r)=r/(r+k) (Monod 1949), where the Monod constant *k* (µg/mL) is the concentration of limiting resource where the growth rates are half their maximum value. The phage adsorb to the sensitive, S, and mucoid, M, cells respectively with rate constants δ_S_ and δ_M_ (hr^-1^mL^-1^) but do not adsorb to the envelope-resistant bacteria N_R_. Upon adsorption, both mucoid and sensitive cells produce β phage particles (neglecting the latent period, for simplicity). Transition from sensitive to mucoid states S→M occurs at a rate *x*, and from mucoid to sensitive M→S at a rate *y*, both per cell per hour with all rates proportional to ψ(r). As in (Chaudhry et al. 2018) we assume that the limiting resource is taken up at a rate proportional to the net growth rate of each bacterial population, modified by the resource conversion parameter *e* (µg), which is the amount of limiting resource needed to produce a new cell (Stewart and Levin 1973). Serial transfer is simulated by diluting the population of bacteria and phage by a factor *d* = 0.01 every 24 hours, removing the existing resource, and adding C µg/ ml of the limiting resource.

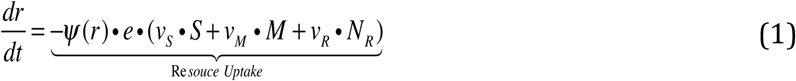

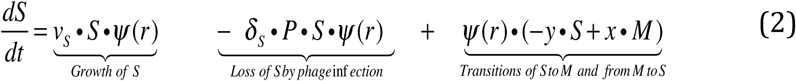

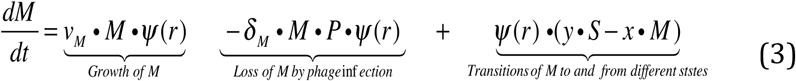

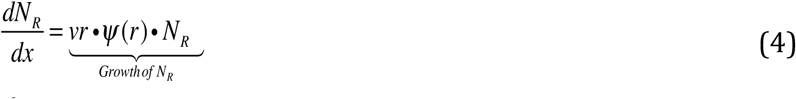

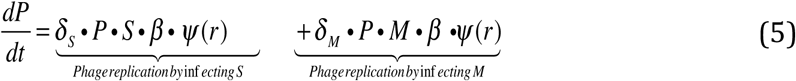

*where* 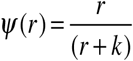

These equations were solved numerically and the dynamics were simulated with Berkeley Madonna™. The values of the phage infection parameters δ_S_ and δ_M_ for the sensitive cells used in these simulations are in the range estimated in (Chaudhry et al. 2018; Pleška et al. 2018) and the growth rates of the bacteria in the ranges estimated in this study (Figure S4). For the mucoids, we postulate the adsorption rates of the phage, which is too low to estimate directly because the rate of decline in phage density due to adsorption to mucoids is within the range of phage deactivation due to other processes in that media. We also postulate different transition rates between the states. Save for the phage λ, we are unable to directly estimate the rate of transition from mucoid to sensitive, M→S, the parameter y. In our analysis, we explore the sensitivity of the dynamics and predictions of the model to the values of the postulated parameters. Copies of the programs and instructions for their use are available from.

### Simulation results

In Figures 5A and 5B, we follow the changes in the densities of the bacterial and phage populations in simulated serial transfer populations initiated with phage and sensitive bacteria, the theoretical analogs of the serial transfer experiments presented in in Figures 1 and 3. In Figure 5 C, D, E and F we follow the changes in the density of bacteria and phage in populations initially dominated by mucoid bacteria; the theoretical analog of the experiments presented in Figures 2. In all of these simulations we are assuming that sensitive cells are the most fit, resistant cells have the next highest fitness, and the mucoid cells have the lowest fitness, respectively v_S_ =2.1, v_NR_=2.0, and v_M_ =1.8. This assumption is supported by our estimates of the maximum growth rates of these different cell lines, and consistent with the evidence present in (Wielgoss et al. 2015) about mucoidy engendering a substantial fitness cost.

**Figure 5:**
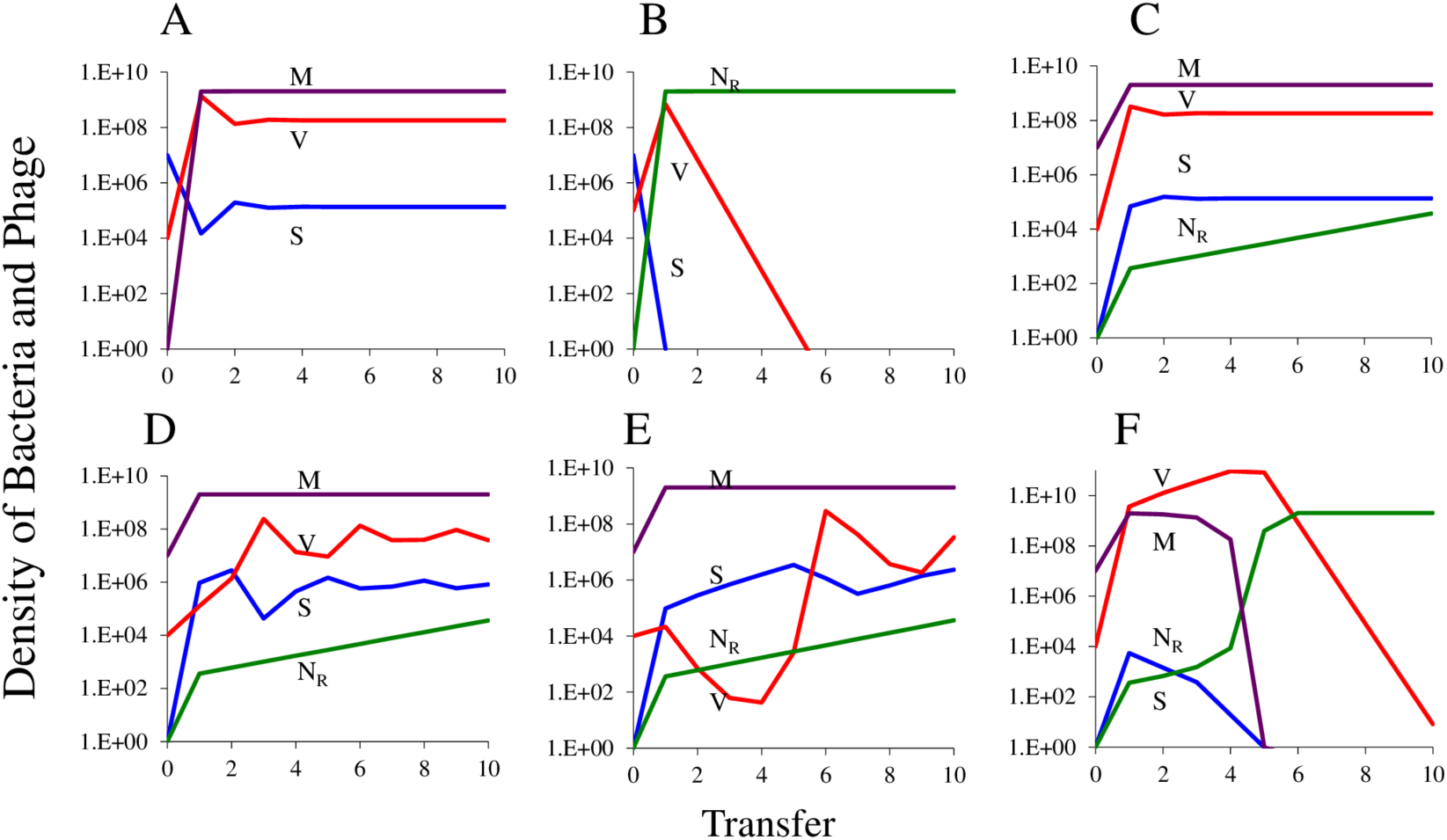
Simulation results of population dynamics in serial transfer experiment of a lytic phage with sensitive, mucoid and resistant bacteria. Changes in the densities of mucoid, M (purple), sensitive, S (blue), and envelope-resistant, N_R_ (green) bacteria and virulent phage, P (red) in populations initially dominated by mucoids. Standard parameters, C=1000, e=5×10^7^, v_S_ =2.1, v_M_ =1.8, and v_R_ =2.0, k=1, β=50, x=10^−5^, δ_S_ = 10^−7^. (A) Establishment of the mucoids in a population of sensitive bacteria and phage in the absence of resistance, y=10^−3^, δ_S_=10^−11^. (B) Establishment of resistant mutants in a population that is unable to generate the mucoids. (C) Establishment of phage in a population dominated by mucoid cells and single sensitive and resistant bacteria, y=10^−3^, δ_S_=10^−11^. (D) Establishment of phage in a population dominated by mucoid cells and single sensitive and resistant bacteria, <u>y=10^−4^</u>, δ_S_=10^−11^. (E) Establishment of phage in a population dominated by mucoid cells and single sensitive and resistant bacteria, <u>y=10^−5^</u>, δ_S_=10^−11^. (F) Establishment of phage in a population dominated by mucoid cells and single sensitive and resistant bacteria, y=10^−3^, and <u>δ_S_=10^−10^</u>.

At the start of the simulations in Figure 5A, there are 10^6^ phage, 10^7^ sensitive cells, a single mucoid cell, no resistant mutants, and the resource is at its maximum level of r=1000. Within a single transfer the mucoid bacteria ascend to become the dominant population of bacteria and by the third transfer the population of bacteria and phage are at equilibrium. If mucoids are not present and cannot be generated (y=0) but resistant mutants are present, they will ascend to become the dominant bacterial strain and the phage will be lost (Figure 5B). Although the dynamics are different than those observed in this simple model, there is qualitative agreement between the theory and experiments. In Figure 1 with T3 and T4, where mucoid bacteria ascended to dominate, the phage were maintained at high densities. In experiments with T5, resistance evolved instead of mucoidy and phage were lost. In one of the three experiments with T7, mucoidy evolved and phage were maintained. In two experiments with T7, neither mucoidy nor resistance evolved; the cells were killed by phage and the phage were washed out in subsequent transfers.

To get a better idea of the conditions required for mucoid bacteria to maintain the phage, and why resistant mutants may not have ascended to dominance in the serial transfer experiments presented in Figure 2, we simulated situations where phage are introduced into populations of mucoid bacteria. At the start of these simulations there are 10^6^ phage, 10^7^ mucoid bacteria, single sensitive and resistant cells (S=1 and N_R_=1), and the resource is present at its maximum level r=1000. The results of these simulations illustrate the critical role that the rate of transition between mucoids and sensitive bacteria, the parameter y, plays in these dynamics. When y =10^−3^, within short order the phage established a stable population in a community dominated by mucoid bacteria (Figure 5C). When y =10^−4^, the phage become established, but their densities oscillate before the population of bacteria and phage becomes stable (Figures 5D and S2A). When y=10^−5^, the oscillations in the densities of the phage are greater (Figure 5E) and stability is not achieved (Figure S2C). Because the resistant mutants in these simulations are more fit than the mucoids, their densities increase at each transfer. However, during the course of these simulations, these resistant mutants may not reach levels where they affect the dynamics of the phage and are unlikely to be detected in the experiments. While this is consistent with some of the experiments where phage are introduced into mucoid populations, as noted in the legend to Figure S1 for both T4 and T5, by the end of the experiments, mutants resistant to these phage ascended and mucoidy declined. If the adsorption rate constant <u>□</u>_<u>S</u>_ is increased to 10^−10^, the phage are not maintained and the higher fitness resistant population ascends to dominate the bacterial population (Figure 5F).

In the simulations presented in Figure 5, two not mutually exclusive mechanisms can be responsible for the maintenance of phage. Both of these mechanisms involve the continuous production of populations of sensitive bacteria of sufficient density to support the replication of the phage population. In the case of resistance, the sensitive population is maintained because they are more fit than the resistant (Levin, Stewart, and Chao 1977; Chao, Levin, and Stewart 1977; Lenski and Levin 1985). In the case of mucoidy, the sensitive cells are maintained because of a continuous transition from the mucoid to sensitive states, the “leaky mechanism”. The dynamics anticipated by these mechanisms are, however, different. This is illustrated in Figure S2.

With the standard parameters and initial conditions used in the 10-transfer simulations presented in Figure 5D, in the longer term the initial oscillations observed in the densities of the phage are damped and the phage are maintained by a minority population of sensitive cells sustained by the fitness cost of resistance, v_R_=2.0 and v_S_=2.1 (Figure S2A). If, however, the parameters are changed so that the phage cannot adsorb to the mucoid cells, □_M_=0 rather than 10^−11^, the oscillations in the densities of sensitive cells and phage continue but do not increase in amplitude (Figure S2B). If the rate of transition from sensitive to mucoid is reduced to y=10^−5^, the amplitude of the oscillations in the densities of phage increases to the point where the phage are lost (Figure S2C). This instability of communities where phage are maintained by the fitness cost of resistance can be seen when the initial density of phage is less than that predicted for the equilibrium (Figure S2D). These oscillations of increasing amplitude do not obtain when the phage are maintained by the transition between mucoid and sensitive cells (Figure S2E). Finally, and most critically, in contrast with leaky resistance in an envelope-resistant population, the leaky resistance mechanism for the maintenance of phage in a mucoid population does not require a fitness cost of the mucoid cells (Figure S2F).

### IV. Test of the leaky resistance hypothesis

In accord with the model, for the sustained density of sensitive cells to be sufficiently great to maintain the phage at the high densities observed, the rate of transition or reversion from mucoid to sensitive non-mucoid bacteria has to be high, on the order of 10^−4^ to 10^−3^ per cell. The phage adsorption rate and fitness cost parameters also have to be in the range modeled for resistant cells not to ascend to dominate the culture and lead to the loss of the phage. More information about the range of parameter values needed to account for the population dynamics observed can be seen in Figure S2. What is clear from this sensitivity analysis is that for mucoids to dominate and the phage to be maintained, the rate parameter of adsorption of the phage to mucoids has to be low, δ_M_<10^−11^. In the following, we present two lines of evidence in support of the hypothesis that there is a high rate of transition from mucoid to sensitive, on the order of 10^−4^ per cell per hour.

### Population dynamic evidence for a high rate of reversion from mucoids to non-mucoids

One line of evidence for a high rate of transition from mucoid to sensitive non-mucoid states comes from population dynamics experiments using a method similar to that employed in (Chaudhry et al. 2018) with genetically marked λ temperate phage, λ^CAM^. This method is based on the premise that lysogens will only be formed in populations of resistant mucoid bacteria by infecting cells in minority population of susceptible non-mucoid bacteria. We first performed experiments demonstrating that like the T-phages, λ^VIR^ maintains high densities in mucoid populations of different types (Fig. S3); notably, as for T5 phage (Fig. 1), mucoidy does not arise in an initially sensitive wild-type population in this medium. We then introduced □^CAM^ to cultures of T3 and T7 phage selected mucoids and the *rcsC*137 constitutive-mucoid mutant. We followed the changes in the densities of free phage, the total densities of cells and, by selective plating (see the Materials and Methods), the lysogens and non-mucoid (sensitive) cells generated during the course of the experiment (Figure 6, A, B, and C). While the rates of accumulation of lysogens varied, these data indicate that reversion from mucoid to non-mucoid occurred frequently during growth for all strains.

**Figure 6:**
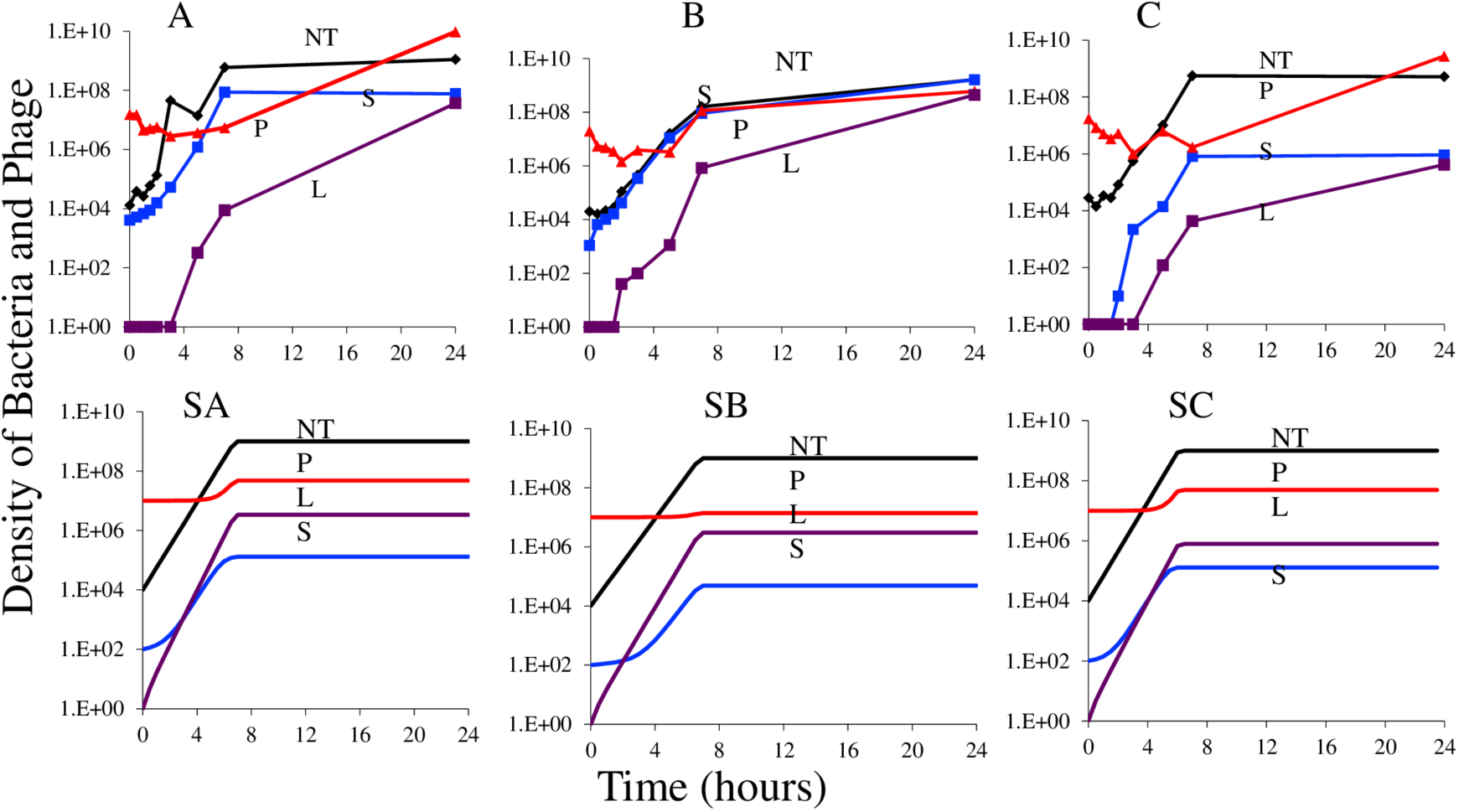
Short-term population dynamics of lysogen formation. A, B and C: experimental results, changes in the densities of total cells NT (black), free λ ^CAM^ phage P (red), lysogens L (purple), and sensitive non-lysogens revertants (blue) S. (A) T3 selected mucoids, (B) T7 selected mucoids, (C) genetically constructed mucoid *E. coli* MG1655 (*rcsC*137 *ompC*::Tn5). SA, SB and SC is simulation results to explain experimental data presented in panel A, B and C with different reversion rate from mucoid to sensitive. SA, SB, and SC: simulations, changes in population density, standard parameters, v_S_=2.1, v_L_=2.1, δ = 2E10^−7^, δ_M_ =10^−11^, x=10^−5^, C=1000, k= 1, e=5×10^−7^, probability of lysogeny, λ= 0.01, i = 10^−4^. (SA) Standard parameters with y=10^−3^, v_M_=1.8, (SB) standard parameters with y=10^−4^ v_M_=1.5., v_M_=1.8, (SC) Standard parameters with y=10^−3^ and v_M_=1.5.

To obtain a rough estimate of the rates of formation of lysogens, we used a simulation similar to that employed in (Chaudhry et al. 2018) (see the supplemental material for the equations and the description of the model) as seen in Figure 6 SA, SB and SC. It is clear that this simple model is not a strictly accurate analogue of the short-term population dynamics of lysogen formation for several reasons. First, in the simulations, □^CAM^ phage does not increase in density at a rate even close to that observed, nor do they reach as high a density. Next, save for the *rcs*137 MUC strain, the predicted density of revertants (S) in simulation is lower than observed in experiments. Most importantly, the predicted rate of increase in the density of lysogens for the T3 and T7 selected mucoids is, with the M > S transition rate of 10^−3^ per hour, less than that observed experimentally and, for the *rcs*137 mucoids, similar to that observed. Nevertheless, these simulations demonstrate that a high rate of transition from mucoidy is necessary to even approximate the observed trends in lysogen formation in an initially mucoid population of cells.

Additional evidence for a high rate of transition from mucoid to non-mucoid and presumably sensitive cells is presented in Figure S4A-D. In this experiment, we followed the numbers of mucoid and non-mucoid colonies in serial transfer culture. It is clear that for the T7 and T4 phage generated mucoids, there is a high rate of transition to non-mucoid. This is not the case for the T3-generated mucoids or the genetically constructed mucoids (*rcsC*137). In Figure S4E we present the estimates of the maximum growth rates of wild-type MG1655, various mucoid lineages, and mutants with envelope resistance against T5 and λ^VIR^. These data indicate that mucoidy does incur a substantial growth rate cost, as indicated by the model analysis. Furthermore, envelope resistance to T5 and λ^VIR^ incurs no such defect under these culture conditions, and may well be the reason why mucoidy is not observed when *E. coli* K12 is challenged with the phagesT5 and λ^VIR^ (Figure 1, Figure S4 E-F). Notably, we observed that these revertants to a non-mucoid phenotype were in some cases able to return to mucoidy upon re-exposure to phage (Figure S5).

### (ii) Single-cell observations of mucoidy, reversion-induced heterogeneity, and phage infection

The second line of evidence for a high rate of transition from mucoid to normal colony sensitive cells comes from single-cell observation of phage infection in mucoid and non-mucoid cultures. Prior work has shown that reversion from phage-induced mucoidy occurs through secondary mutations within the Rcs system rather than through direct reversion of the initial mutation (Wielgoss et al. 2015), and that these mutations restore expression of Rcs to roughly wild-type levels. If this is the case, it should be possible to identify individual mucoid cells based on expression levels in the Rcs pathway. To this end, we used a plasmid encoding a fluorescent reporter for Rcs expression (pBAD18Cm::P_rprA_-mCherry) to visualize heterogeneity within initially mucoid populations (Figure 7, Figure S4, Figure S6). Mucoid and non-mucoid cells do not appear morphologically distinct on a single-cell level, and mucoids do not clump or agglomerate in well-mixed liquid culture, which might have protected these cells against phage (Vidakovic et al. 2018). However, fluorescence expression from the Rcs promoter is clearly different between wild type and mucoid lineages.

**Figure 7:**
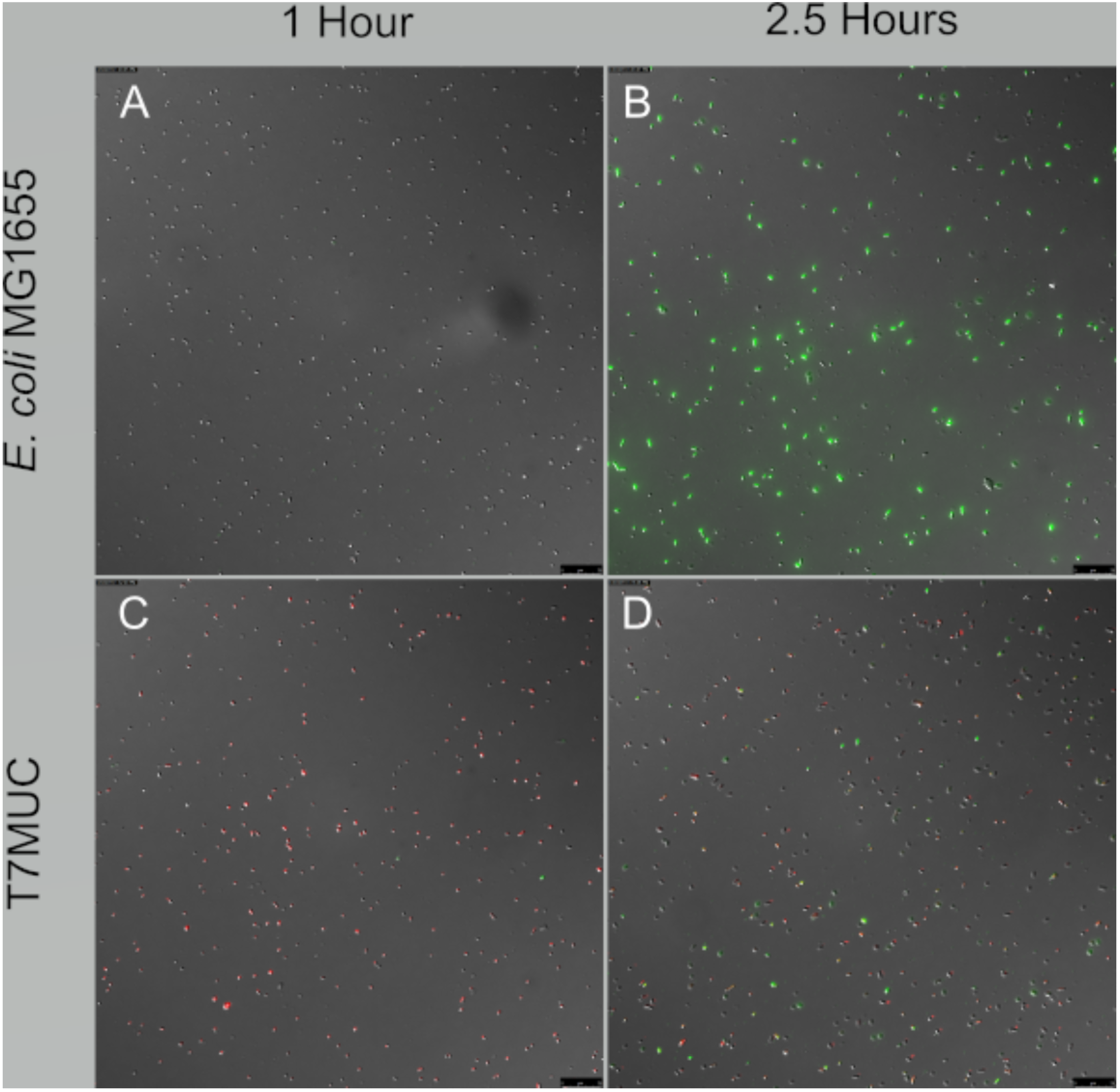
Rcs pathway induction is negatively correlated with T7 phage infection on the single-cell level. In MG1655 wild-type non-mucoid culture (top), expression of Rcs (red) is low, and T7 phage infection (green) is universal. By contrast, in T7-induced mucoid culture (bottom), Rcs expression is high in the bulk of the population and low in a minority of putative revertants, and only the low-Rcs sub-population is infected with T7 phage. Images represent separate aliquots from 1 mL LB cultures incubated at 37°C, sampled approximately 1 hour (left panels) and 2.5 hours (right panels) after inoculation. Images are an overlay of DIC, TRITC, and GFP channels, acquired using a 40X objective (total magnification 400X). Scale bar is 25 µm.

If reversion to non-mucoid eliminates over-expression of Rcs and consequently increases sorption by virulent phage, this should be visible as differences in single-cell sorption to phage which anti-correlate with expression of the Rcs pathway. Using cells expressing the mCherry Rcs fluorescent reporter as described above, we infected cultures with GFP-fluorescent T7 phage (Vidakovic et al. 2018). The T7-derived mucoid lineage was chosen due to the rapid reversion to non-mucoidy and high expected levels of heterogeneity in these cultures, and wild-type non-mucoid *E. coli* MG1655 was used for comparison. We observed that, in wild-type cultures of *E. coli* MG1655 where expression of Rcs is uniformly low, most to all individual cells adsorb phage and become infected (Figure 7A,B; Figure S6). By contrast, in T7-induced mucoid *E. coli*, the majority population of cells express Rcs much more strongly than the non-mucoid MG1655 cells (Mann-Whitney U test on RFP fluorescence at 2.5h, n1=572, n2=434, W=35225, p<2.2e^-16^) do not become infected with the phage (Mann-Whitney U test on GFP fluorescence at 2.5h, n1=572, n2=434, W=35225, p<2.2e^-16^) (Fig. 7 C, D, S6 Fig). However, a minority population of low-Rcs cells, apparently corresponding to the revertants expected in this culture, become infected with T7 phage (see Figure S6).

### (iii) Molecular genetics of mucoid to non-mucoid reversion

The nature of the mutations that are responsible for transition mucoid to phage-sensitive non-mucoid colony also supports a model of high transition rates. Mucoidy results from gain-of-function mutations that strongly activate the Rcs signaling and hence colanic acid production. Non-mucoid reversion can occur via loss-of-function mutations in any of at least six genes (*rcsABCD* and *cpsBG*). Using whole-genome sequencing, we identified the genetic basis for mucoidy and non-mucoidy reversion from T3, T4, and T7 selected colonies (Table S2). Notably, we found instances of IS element insertion in the *rcs* genes. IS elements are abundant in E. coli K12 (seven copies each of IS*1* and IS*2*, and eleven copies of IS*5*, among others). IS elements are able to transpose at relatively high frequencies estimated between 10^−3^ and 10^−6^ per element per generation (Craig 1996). In all, molecular genetics offers a basis for high frequency of reversion to non-mucoid: (i) there is a large set of target genes, any of which can be inactivated to yield revertants; and (ii) there is an abundance of chromosomal IS elements that frequently transpose and can inactive genes. Hence, mutagenesis via IS element insertion can occur at much higher rates than spontaneous nucleotide changes and in the order of frequency we propose.

## Discussion

A longstanding, fundamental question about the ecology, population and evolutionary biology of bacteria and phage is how are these viruses maintained in bacterial communities following the ascent and dominance of populations of bacteria upon which they cannot replicate (Campbell 1961; Levin, Stewart, and Chao 1977; Chao, Levin, and Stewart 1977; Gomez and Buckling 2011; Wei, Kirby, and Levin 2011; Jover, Cortez, and Weitz 2013; Jover, Romberg, and Weitz 2016; Weissman et al. 2017)? The results of this experimental study with *E. coli* K12 and five of its lytic phage, T3, T4, T5, T7 and □^VIR^, provide a new and general answer to this question in the form of a long recognized mechanism of phage resistance, mucoidy. We present evidence that when serial transfer populations of *E. coli* are confronted with some but not all of these lytic phage, mucoid colony producing bacteria ascend and become the dominant population of bacteria. The phage continue to be maintained for extensive periods at high densities with the dominant mucoid population of bacteria at densities not much less than that of phage-free cultures with those bacteria.

As anticipated from the study by Wielgloss and colleagues (Wielgoss et al. 2015), the resistance provided by mucoidy is general; mucoid colony producing bacteria are resistant to phages with different receptor sites (Wright et al. 2018). Not presented in that or earlier investigations, is the observation presented here that phage can be maintained in these mucoid populations of bacteria, or a mechanism that can account for this maintenance. Moreover, the results presented here indicate that whether the phage select for the mucoid bacteria they are cultured with or not, or whether these mucoid colony-producing *E. coli* are the product of phage-mediated selection or genetic manipulation, all five lytic phage studied maintain stable populations with these mucoid bacteria.

Based on the results of our analysis of the properties of a mathematical model, we postulate that a sufficient, if not the unique, mechanism by which the phage are maintained in these mucoid populations of *E. coli* is a high rate of transition from the resistant, mucoid state to the susceptible state, “leaky resistance” (Delbruck 1946; Weissman et al. 2017; Chaudhry et al. 2018). We cannot, however, rule out some contribution of the fitness cost of mucoidy contributing to the maintenance of the phage (Levin, Stewart, and Chao 1977; Lenski and Levin 1985). Central to the leaky resistance hypothesis is that the rate of transition from the mucoid to the susceptible state is high, on the order of 10^−4^ to 10^−3^ per cell per hour.

*A priori*, a transition rate between phenotypic states of bacteria of this great a magnitude by classical genetic processes is highly unlikely. To be sure, the large size of the Rcs regulon provides an abundance of mutational targets (Wielgoss et al. 2015). Nevertheless, *E. coli* is only expected to experience roughly one mutation per thousand cells per generation overall (Lee et al. 2012), which, no matter how big the target may be, means that a rate of 10^−4^ is too high to explain by base substitution mutation. However, the number of transposable IS elements encoded in E. coli and their high rate of transposition offer another mechanism by which to inactivate *rcs* or *cps* genes. Indeed, we have direct evidence that IS elements are responsible for some of our non-mucoid revertants.

Consistent with other studies, we found that some mucoid phage-resistant mutants revert quickly to a non-mucoid, phage-sensitive state when the phage are removed, due to mutations elsewhere in the Rcs pathway rather than through reversion of the original mucoidy-inducing mutation (Wielgoss et al. 2015). Reversion appeared to occur at different rates in *E. coli* mucoids obtained through exposure to different T phage, such that T3-derived mucoids appeared to be relatively stable and T4- and T7-derived mucoids reverted more rapidly. Observations of the growth rate defect in mucoid cells (Figure S5.) indicated that T3-derived mucoids had a strong growth defect, suggesting that the difference in observed reversion was not due to a lack of selective pressure for improved growth. It may therefore be the case that the specific mutation(s) that produce mucoidy in our T3-derived lineages are more difficult to reverse than the mutations that arise in other phage-selected mucoid lineages, or that direct reversion of the original mutation is required.

Also consistent with the postulated high transition from mucoid to susceptible are the results of the experiments we have done to estimate rate of production of lysogens for the phage □. When genetically marked temperate □ are mixed with □ resistant mucoids populations generated in different ways, the rate of production of lysogens is consistent with the transition rate from mucoid to susceptible being on the order of 10^−4^ per cell per hour. While not providing estimates of the transition rate, additional support for this rate being high, comes from the experiments we did with single-cell microscopy. We demonstrate that propagation of phage in cultures of mucoid *E. coli* relies on infection and lysis of these revertant cells, which have restored wild-type levels of expression through the Rcs capsule synthesis system. Our sequencing data rationalize how high frequency of reversion can occur at the molecular genetic level. We detect IS elements disrupting *rcs* genes to counter the effect of mutations that activate Rcs and induce mucoidy. Owing to their high frequencies of transposition, IS elements disruption of the rcs genes offer a ready explanation for the high frequency transition to non-mucoid revertant.

From the sequence data, can we argue that the postulated high rates of reversion from mucoid to sensitive, can be attributed to what can be broadly described as “genomic instability”. As a consequence of amplification of regions of the chromosome (Nicoloff et al. 2019; Andersson, Nicoloff, and Hjort 2019; Anderson et al. 2018), unstable chromosome arrangements (Guerillot et al. 2019), and movements of insertion sequences (Chaudhry et al. 2018), seeming genetically homogeneous populations of bacteria include minority populations that are genetically and phenotypically different than the majority. The rates of transition to and from these unstable modifications to the stable state can be quite high, 10^−3^ per cell per hour or more. When selected for, these minority populations ascend, which is what we postulate is happening with the generation resistant mucoid bacteria when they are confronted with phage. When these minority populations are no longer favored, in the absence of phage in the muco*id E. coli*, or in the absences of antibiotic for heteroresistance (Nicoloff et al. 2019; Andersson, Nicoloff, and Hjort 2019; Anderson et al. 2018), the populations revert back to their ancestral state.

We observed that, in some cases, susceptible revertants from mucoidy were able to regain the mucoid phenotype upon re-exposure to phage. This is in contrast with a previous study which found that mutations leading to reversion of mucoidy prevented a return to the mucoid phenotype (Wielgoss et al. 2015). In that study, most of the tested non-mucoid revertants had acquired large gene deletions spanning the *rcs* operon. The nature of these mutations likely explains why subsequent return to the mucoid phenotype was not possible: deletions are a loss of genetic information that cannot be reverted. We therefore suggest that reversion back to mucoidy occurs via other means in our study. These might be nucleotide changes that cause gene loss-of-function that can subsequently be reverted to wild type. Alternatively, these might be IS element excision when these mobile genetic elements are either spontaneously excised or moved via a cut-and-paste mechanism.

While we have focused on the role of resistance generated by mucoidy, the results of our experiments do not rule out a substantial role of surface, or envelope, resistance in the dynamics of *E. coli* and its lytic phage. Indeed, if envelope resistant mutants are less costly than mucoid, any envelope resistant mutants that do enter a community of bacteria and phage will increase in frequency. Consistent with this prediction is that in some cases we observed envelope resistance arising within mucoid cultures. Most notably this occurs with T5 phage, where envelope resistance is expected to have minimal or no cost (Lenski and Levin 1985). There is also evidence that mucoidy and other resistance mechanisms can arise together within single cultures (Wielgoss et al. 2015). Our results indicate that the evolution and utility of mucoidy, as compared with other phage resistance mechanisms, will depend heavily on the strains involved and the environmental context of the interaction. Under these conditions (laboratory *E. coli* exposed to single strains of phage in liquid media), the exponential growth rate was a reasonably informative measure of fitness, but this is known to be untrue in other scenarios (Concepción-Acevedo et al. 2015), and particularly in more complex environments (Gomez and Buckling 2011; Bjorkman et al. 2000; Santos-Lopez et al. 2019). Understanding the selective forces that are likely to dominate in different contexts will therefore be relevant for a global understanding of mucoidy as a part of phage-bacteria interactions. Finally, our models and experimental results suggest that if mucoidy evolves first, mutants with more specific resistance mechanisms are, with the exceptions like T5, unlikely to ascend.

### Caveats

At one level, the population dynamic and experimental and modeling results presented here are mechanistic, in that they explain how mucoid *E. coli* can maintain high-density populations of lytic phage. However, as suggested in the preceding, at the genetic, physiological and molecular levels, these results and explanations raise mechanistic questions, which we allude to, but do not fully answer. Included among those unanswered or insufficiently questions are: (1) How does mucoidy provide the broad “resistance” to phages? (2) What might the consequences of mucoidy be for microbial populations in natural environments?

These studies were performed in well-mixed liquid media in serial transfer culture. Under these conditions, wall populations are unlikely to provide a refuge for stationary phase cells and allow the maintenance of the phage in this way (Schrag and Mittler 1996). Moreover, under these conditions multicell collectives are not anticipated to be formed and were not observed. In stationary cultures where surface adhesion and biofilm formation occur, these collectives could allow exopolysaccharide production to be used as a common good (Vidakovic et al. 2018; Malhotra et al. 2018). Bacteria-phage dynamics in surface-adhered conditions might therefore differ from those observed here; for example, creation of surplus extracellular matrix by mucoid bacteria might provide some protection to phage-sensitive bacteria, either by impeding free diffusion of phage or by providing a scaffolding of phage-sorbing cellular material, allowing maintenance of a sensitive sub-population alongside mucoids and phage (Vidakovic et al. 2018). As the Rcs pathway in *E. coli* is a known regulator of biofilm formation (Clarke 2010), it is very plausible that mutations in this pathway that generate mucoidy in the interests of phage resistance will also produce changes in collective behavior.

### A practical implication

While of interest to understanding the how phage are maintained in natural populations of the *E. coli* and role (or lack of role) of these viruses in regulating the densities of these bacteria and determining the strain, we expect the results of these studies with *E. coli* K12 and its phages will be of little clinical or other practical importance. On the other hand, they may well be relevant for understanding the evolution and maintenance of mucoidy in other more medically or agriculturally relevant systems. Mucoidy has been observed in chronic *Pseudomonas aeruginosa* infections of the cystic fibrosis lung and marks progression of the disease into a more debilitating stage (Malhotra et al. 2018; Li et al. 2005). While it is at present not known whether phage-imposed selection plays a role in this clinical phenomenon, selection for mucoidy has been observed in *Pseudomonas* exposed to lytic phage (Martin 1973; Scanlan and Buckling 2012). As interest in phage-based therapy for cystic fibrosis lung infection grows (Cafora et al. 2019; Forti et al. 2018; Alemayehu et al. 2012; Debarbieux et al. 2010), clarifying the role of phage in disease-associated mucoidy will be critical in developing safe and useful treatments for these intransigent infections.

## Acknowledgements

The authors would like to thank Cary Nadell, Jim Bull, and Sylvain Moineau for generously providing us with the phage used in these experiments and Ole Skovgaard for providing the *E. coli* MG1655 employed here. This research was funded by grants from the U.S. National Institutes of General Medical Science GM091875-17 and 1R35 GM136407-01 to B.R.L., 1R35GM133509-01 to M.G., and Emory University start-up funds (MG, NV). We are grateful to Ingrid McCall for managing the logistics of this enterprise and for providing editorial and other suggestions, and to Melony Ivey for superb technical assistance.

## SUPPLEMENTAL MATERIAL

**Figure S1:**
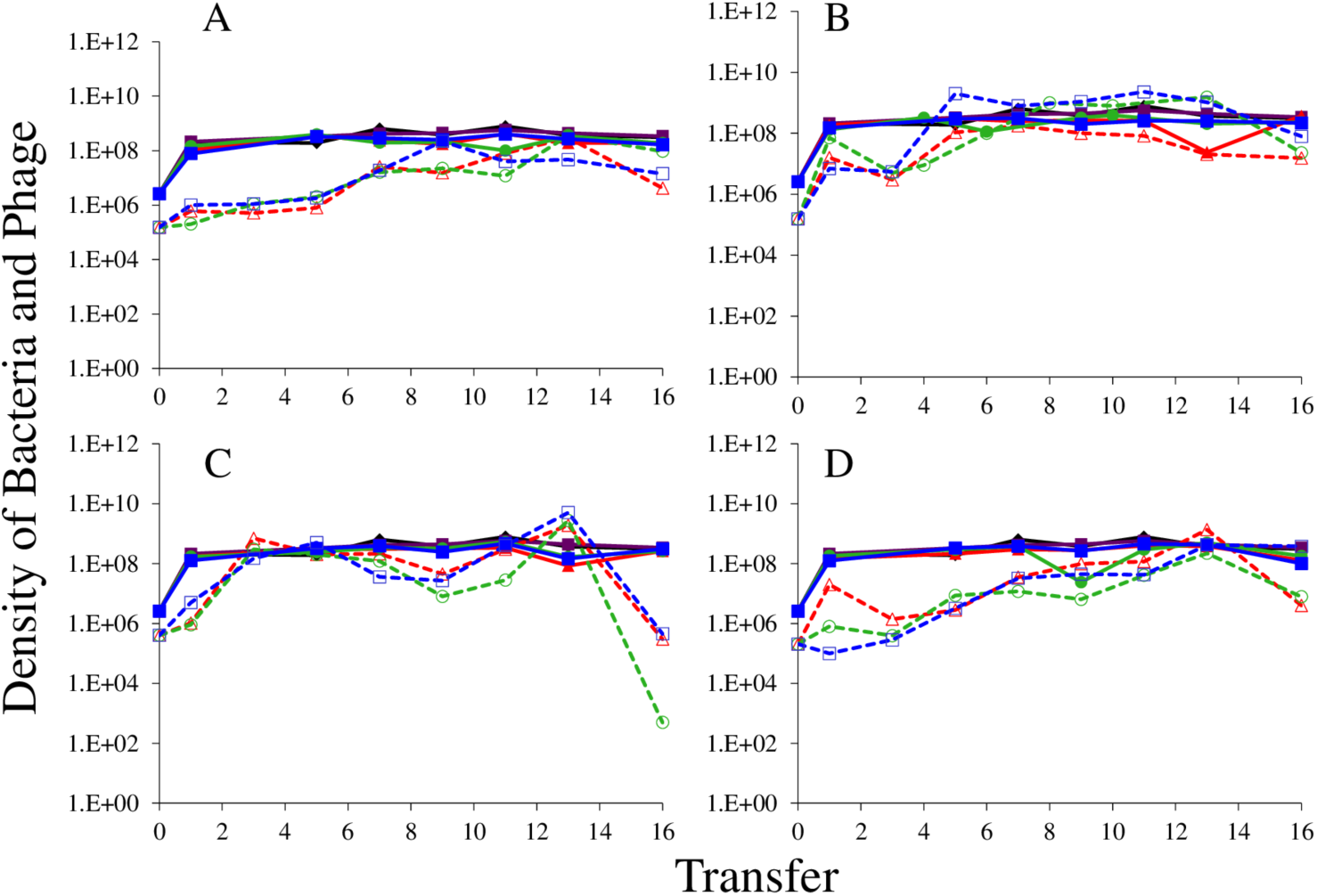
The population dynamics of T3 (A), T4 (B), T5 (C) and T7 (D) phage with T7 generated mucoid *E. coli* MG1655 as the host strain. Cultures were diluted 1:100 into fresh M9 + 0.4% glucose every 24 hours, and CFU-PFU/mL measurements were taken via dilution plating. Solid lines show bacterial densities in each replicate (red/green/blue); dotted lines show the phage densities in the corresponding cultures. Phage-free cultures (black and purple lines) are shown as controls, to indicate resource-limited population density for the bacteria. The phage-free controls lost the mucoid phenotype. At the end of the last serial transfer, three colonies from each phage-infected culture were streaked at least three times to get rid of co-evolving phage. These isolated clones were tested against the phage used in the experiment; envelope resistance arose in T4 and T5-infected cultures, accompanied by a loss of mucoidy.

**Figure S2:**
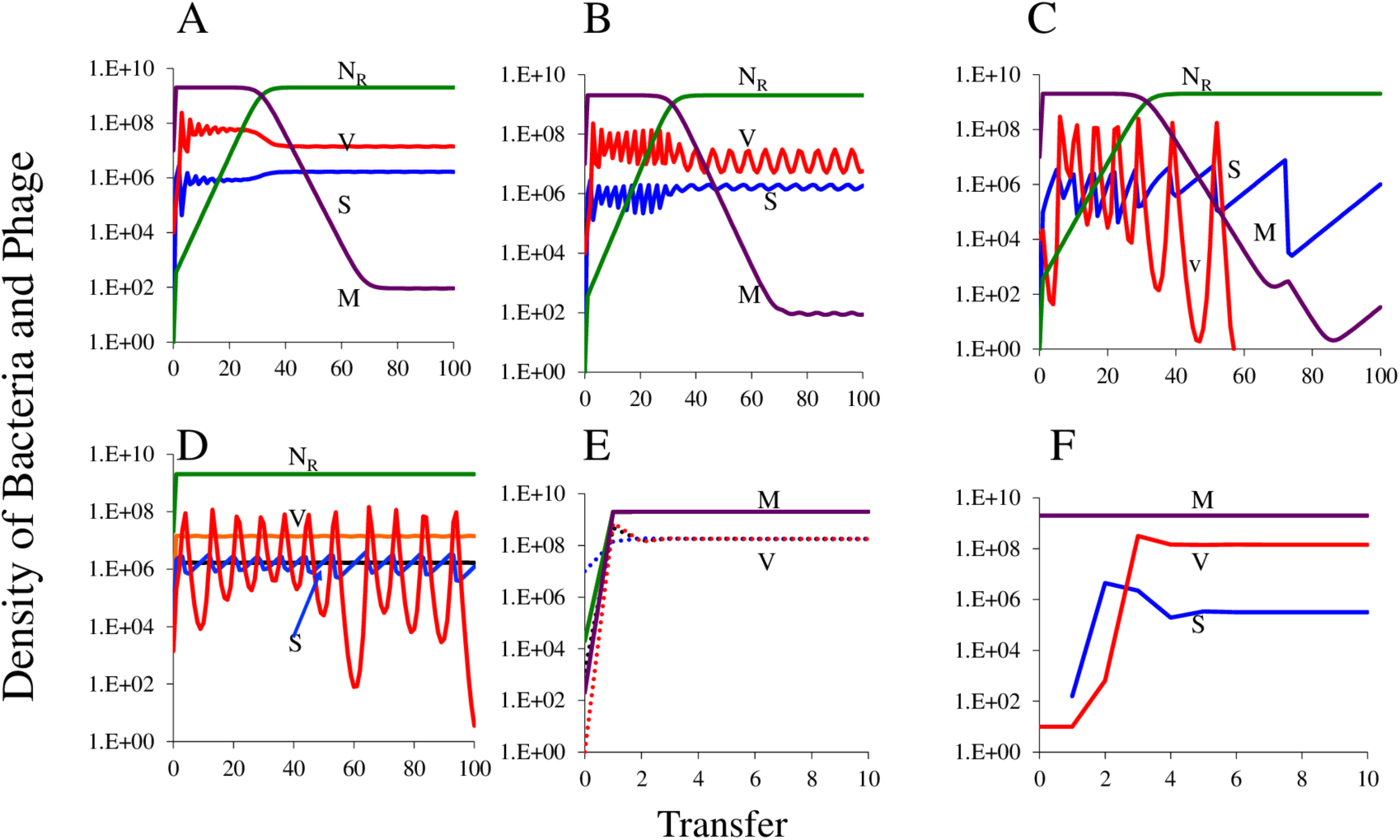
Changes in the densities of bacteria and phage in serial transfer culture. Standard parameters, C=1000, e=5×10^7^, v_S_ =2.1, v_R_ =2.0, k=1, β=50, x=10^−5^, δ_S_ = 10^−7^. (A) Long term dynamics of a population with mucoid, sensitive, and resistant bacteria v_M_=1.8, δ_M_=10^−11^, y=10^−3^, continuation of the dynamics in Figure 5B. (B) Long term dynamics of a population with mucoid, sensitive and resistant bacteria with the mucoid bacteria fully resistant to the phage, <u>δ_M_=0</u>, v_M_=1.8, y=10^−3^. (C) Long term dynamics of a population with mucoid, sensitive, and resistant bacteria with a lower rate of transition from S to M, <u>y=10^−5^</u>, δ_M_=10^−11^, δ_S_=10^−7^, a continuation of the dynamics presented in Figure 5C. (D) Long term dynamics of a community where phage are maintained by the fitness cost of resistance and no mucoid bacteria can be generated. Two simulations, one where the initial densities of phage are the 0.01 of the equilibrium, N_R_=2.0X10^7^, P=1.40X10^5^, S=1.67×10^4^ (see Figure S2A) and one where the initial density of phage is 1.4X10^3^. (E) Dynamics of a community where the phage are maintained by the transitions of mucoid to sensitive (leaky resistance) v_M_=1.8, δ_M_=10^−11^, y=10^−3^, no resistant bacteria; effect of the initial densities of mucoid bacteria and phage. (F) Dynamics of a community where the phage are maintained by the transitions from sensitive to mucoid, but there is no fitness cost of mucoidy, <u>v_M_=v_S_ =2.1</u>, δ_M_=10^−11^, y=10^−3^, δ_S_ =10^−7^, no resistant cells are present or can be generated.

**Figure S3:**
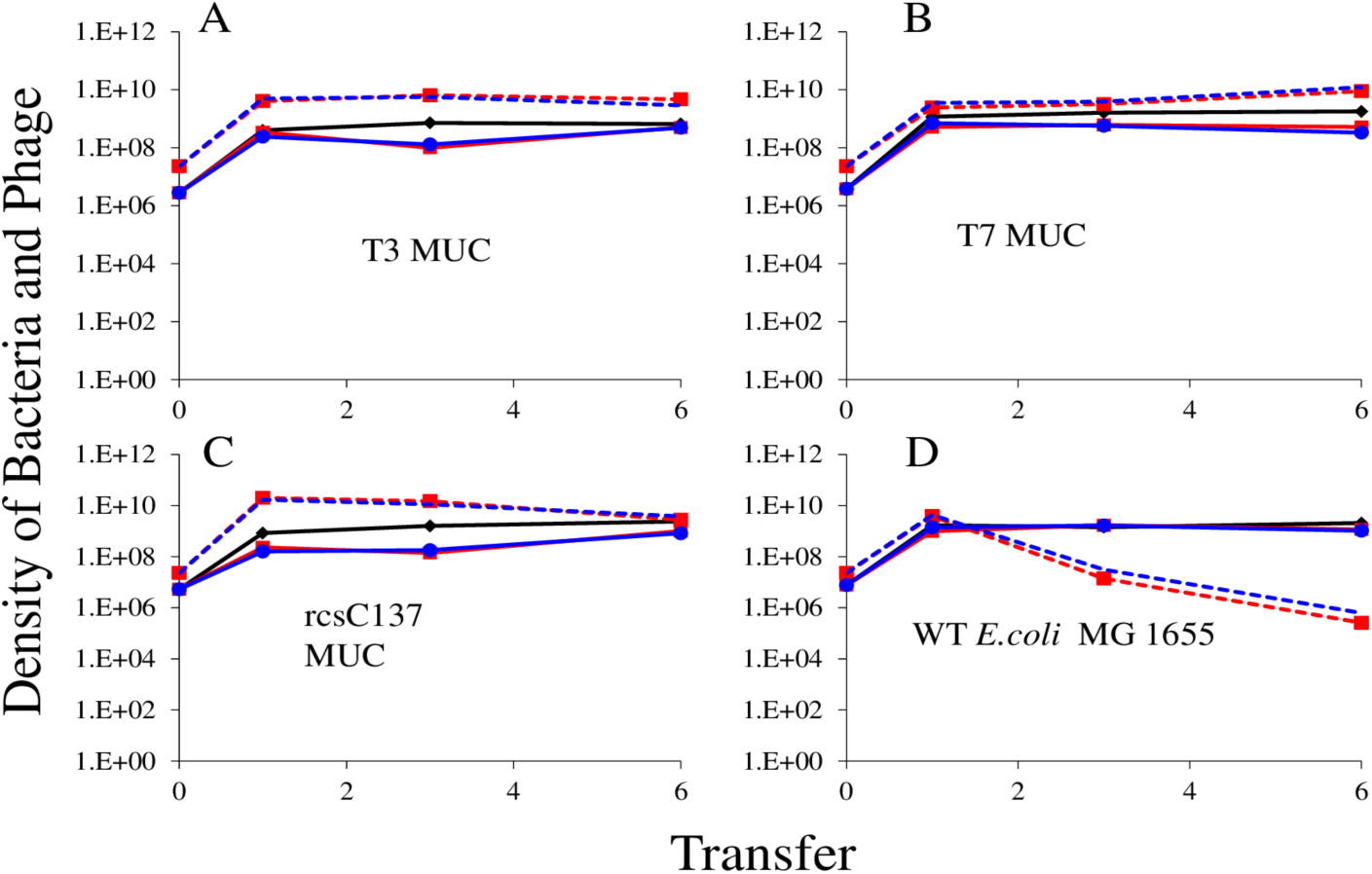
Population dynamics of λ^VIR^ phage with T3-generated mucoid, T7-generated mucoid, *rcsC*137 *ompC*::Tn5 (constitutive mucoid) and WT *E.coli* MG1655. Cultures were inoculated into LB medium with λ^VIR^ phage (1:1) and diluted 1:100 into fresh LB medium every 24 hours; two replicates were performed per condition. The decline of phage in wild-type cultures corresponds to the emergence of envelope resistance against λ^VIR^. Solid red and blue lines show bacterial densities while dotted line show the phage densities in the corresponding cultures. Phage-free cultures (black lines) are shown as controls, to indicate resource-limited population size for the bacteria.

**Figure S4:**
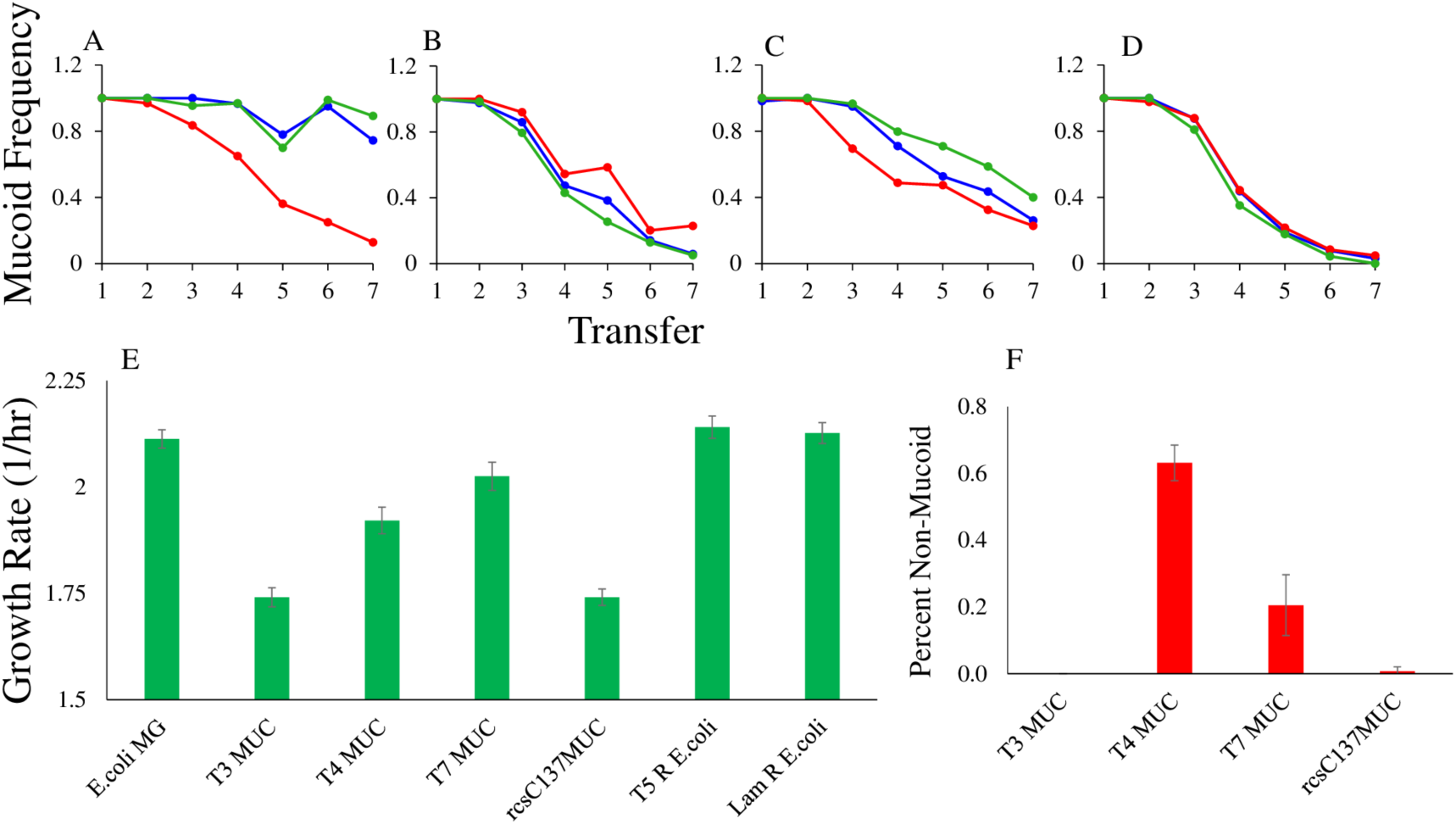
Reversion from mucoid and the growth rates of the different mucoid cell lines. To estimate the reversion rate of mucoid to non-mucoid phenotype, single mucoid colonies were picked from the 9^th^ serial transfer of (A) T3, (B) T4, (C) T7 phage infected cultures of *E. coli* MG1655 (Figure 1) and (D) genetically constructed mucoid *E. coli* MG1655 (*rcsC*137 *ompC*::Tn5) and grown in liquid culture in the absence of phage, with dilution 1:100 into fresh LB broth every 24 hours. At the end of every transfer, the culture was serially diluted and plated on LB agar for CFU/ml counts and colony morphology determination. Red, green and blue lanes represent three independent replicates (E) Mucoid (MUC) cultures suffer a substantial growth rate defect as compared with wild-type *E. coli* MG1655, while envelope resistance (R) to T5 and λ phage incurs no growth rate penalty under these conditions. Note that the “T4 Muc” and “T7 Muc” cultures immediately produce a mixed culture of mucoid and revertant colonies, as seen during CFU plating of 24-hour growth curve cultures (F), and the estimated growth rate for the mixed culture is likely to reflect the more rapid growth rate of the substantial revertant fraction. Data were obtained from OD_600_ growth curves in LB at 37°C. Error bars represent mean ± SD of ten (E) or three (F) replicates.

**Figure S5:**
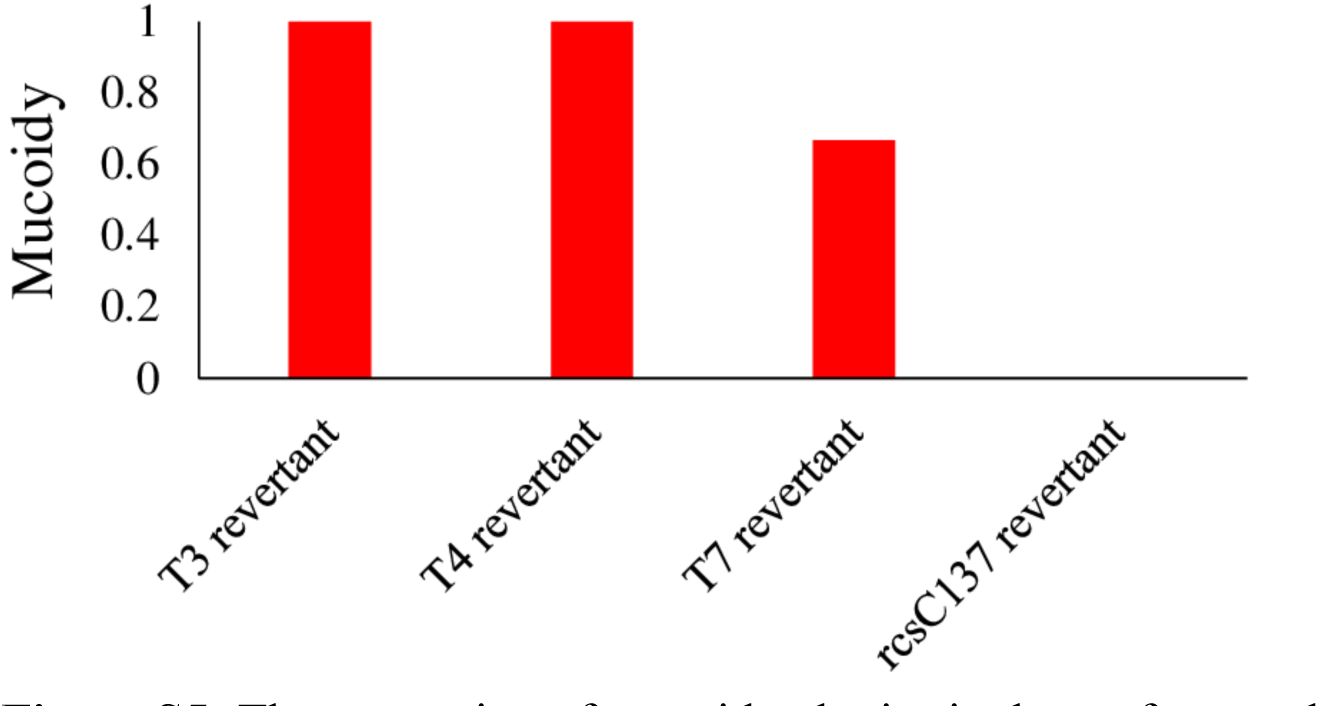
The proportion of mucoid colonies is shown for populations of the ancestral revertant. Mucoidy was scored after exposure to phages on the phage plate. Three individual revertant non-mucoid colonies were picked from each ancestor mucoid background (bacteria from transfer 8 derived from mucoid ancestors, Figure S4 A, B, C, D). 100 ul of overnight cultures of these clones was mixed with 100 µl of a high titer of each respective phage along with 3ml soft agar and poured on the plate; rcsC137 revertants were exposed to T7 phage.

**Figure S6:**
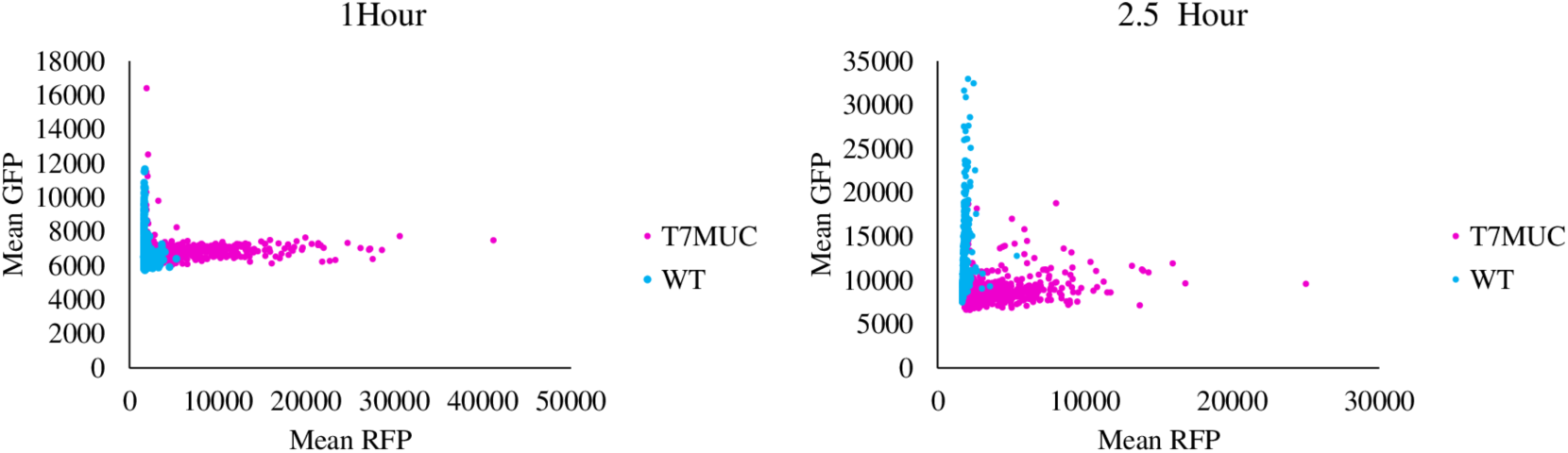
Quantification of red (TRITC) and green (GFP) channels for the images shown in Figure 7. Points represent mean pixel intensity for each channel for objects identified as particles in DIC channel. Image quantification was performed in ImageJ.

**Table S1.** Statistical comparison and quantile values of red (TRITC) and green (GFP) channels for the images shown in Figures 7 and S6, using the Mann-Whitney U test.

|  | Mann-Whitney U Test |  |  |  |
| --- | --- | --- | --- | --- |
|  | T7 vs MG1655, 1h |  | T7 vs MG1655, 2.5h |  |
|  | GFP | RFP | GFP | RFP |
| q25 T7 | 6654 | 2084 | 7957 | 2346 |
| q50 T7 | 6909 | 5088 | 8442 | 3029 |
| q75 T7 | 7128 | 9279 | 9066 | 4669 |
| q25 MG1655 | 6273 | 1700 | 9525 | 1705 |
| q50 MG1655 | 6472 | 1814 | 10650 | 1767 |
| q75 MG1655 | 6785 | 2100 | 13519 | 1894 |
| n1 | 473 | 473 | 572 | 572 |
| n2 | 452 | 452 | 434 | 434 |
| W | 155359 | 189751 | 35223 | 241559 |
| p | < 2.2E-16 | < 2.2E-16 | < 2.2E-16 | < 2.2E-16 |

**Table S2.**
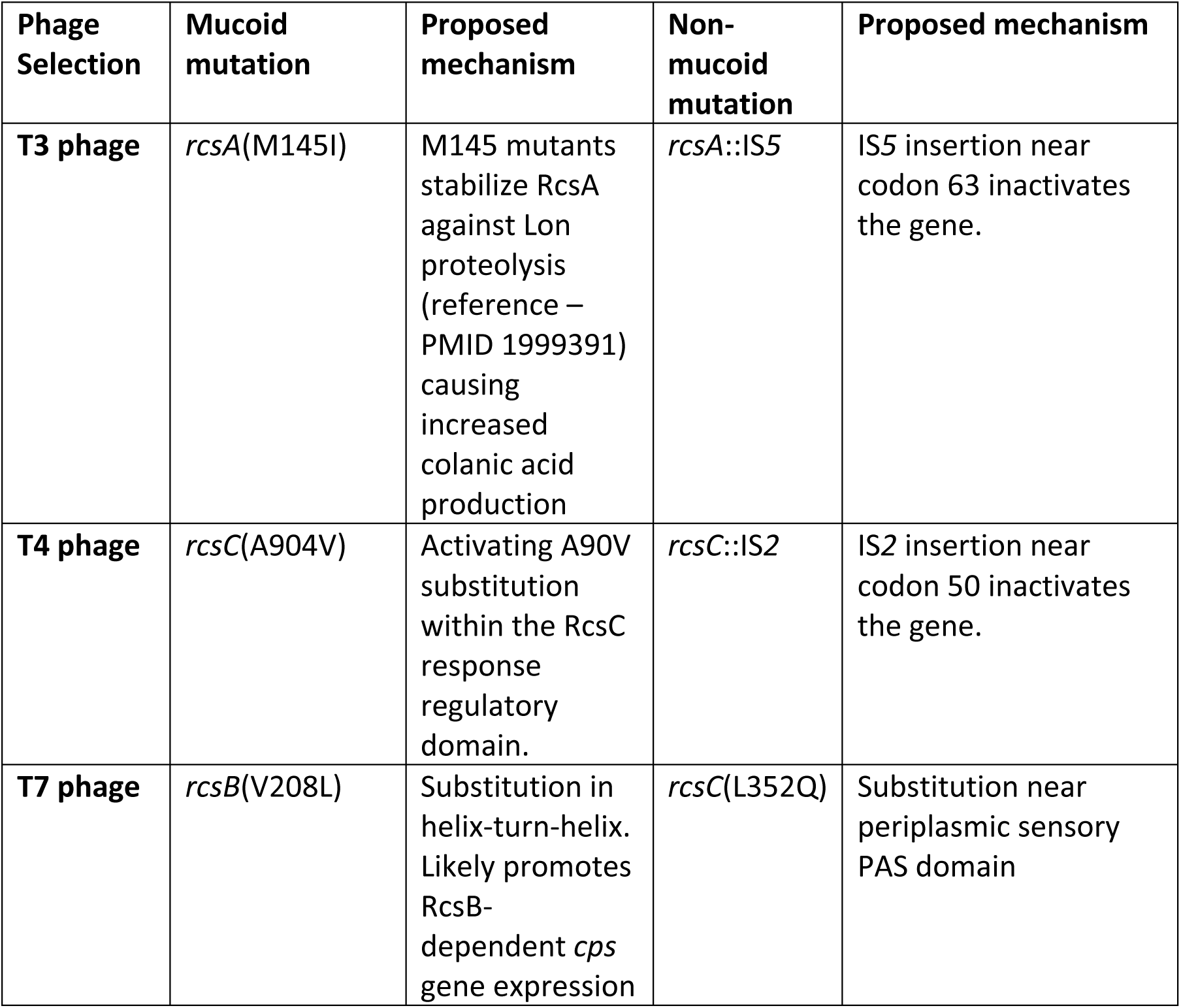
Genetic basis for mucoid mutants and their non-mucoid revertants.

### Model of the dynamics of lysogen formation with leaky resistance

There are three populations of bacteria sensitive non-lysogens, S, resistant (Mucoid), M non-lysogens, and lysogens, L and one population of phage, P, where S, M, L and P are the densities, cells and particles per ml as well as the designations of these populations. The bacteria grow at maximum rates, v_S_, v_M_, and v_L_ with the realized rate of growth proportional to the concentration of the limiting resource,

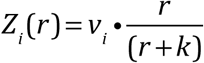

r µg/ml, where i is S, M or L, and k is the concentration of the resource where the realized growth rate is half its maximum value. For the production of a new cell, e µg of the resource has to be consumed.

The phage adsorb to the bacteria at a rate proportional to the product of their densities and a rate constant δ. A fraction, λ (0 < λ <1) of the adsorptions of the temperate phage to sensitive cells, P to S, results in the production of lysogens. Phage that adsorb to lysogens are lost. We assume that the mucoid population is completely resistant to the phage. The (1-λ) fraction of infections of sensitive cells by temperate phage result in the production of β, the burst size, of free phage. With a probability i (0 < i < 1) per cell per hour lysogens are induced and produce β phage. With a rate y per cell per hour, the mucoid, M produce sensitive cells, M→S, and x per cell per hour, the sensitive cells generate mucoids, S→M. With these definitions and assumptions, the rate of change in densities of the bacteria and phage and concentration of the limiting resource are given by the following set of coupled differential equations.

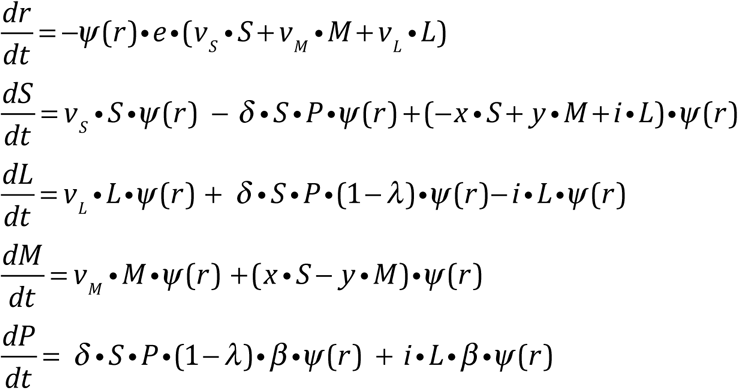

*where* 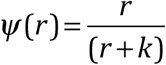

## Notes

### Competing Interest Statement

The authors have declared no competing interest.

